# Species Delimitation Under Allopatry: Genomic Divergences Within and Across Continents in Lepidoptera

**DOI:** 10.1101/2023.03.06.531242

**Authors:** Mukta Joshi, Marianne Espeland, Peter Huemer, Jeremy deWaard, Marko Mutanen

## Abstract

Delimitation of allopatric populations into species remains subjective and largely arbitrary. Many cold-adapted species from the subarctic and Central and Southern European Mountain systems display frequent allopatry with disjunct distributions of their populations. The same concerns Holarctic species which are many and which almost regularly show various degrees of differentiation between the continents. In this study, we analyze high-throughput target enrichment data for ten groups of arctic-alpine and Holarctic lepidopteran species sampled from four main regions across the Holarctic realm – Fennoscandia, North America, Alps and Altai. We first aimed to assess whether the genetic differences in the nuclear genome reflect observed DNA barcode divergences and second, whether the gap between population and species-level differences can be dissected using genomic data. We compared the phylogenetic trees and uncorrected pairwise genetic distances obtained from target enrichment and the mitochondrial cytochrome oxidase I (COI) barcodes for each of the study species. Additionally, we also performed a suite of population genetic and species delimitation analyses to further shed light on patterns of intraspecific variation using a large number of nuclear markers. We observed that in about one half of the cases, DNA barcodes tended to show phylogenetic relationships similar to the target enrichment markers. We report varying levels of nuclear genetic differentiation among the populations analyzed, starting from low differentiation of geographically separated populations to the deeper separation of some Nearctic population and further arctic-alpine disjunction. Given that no single consistent pattern emerged across different case studies, we demonstrate that the delimitation of allopatric populations into species could be done much more efficiently and in a consistent manner if based on a large set of universal genetic loci, which would help in reaching standards for taxonomic delimitation of allopatric populations.

---

Species delimitation, despite being one of the main commissions of taxonomic research, remains challenging. This is largely due to the ontological issues around the term ‘species’ as a variety of different species concepts have been proposed, and there are difficulties in reaching a consensus over them (De Queiroz 2007). Perhaps even more importantly, delimitation is difficult due to obscure boundaries between species under many settings and due to arbitrary boundaries during the long, continuous process of speciation (Nosil, 2008). The traditional way of delimiting species using morphology has limitations, particularly in cases of cryptic diversity, i.e., when the taxa are genetically distinct with only minor or no morphological differences (Knowlton 1993; Packer et al. 2009). Additionally, morphological practices tend to be slow and time-consuming, and are often biased due to personal preferences of the taxonomists (Godfray 2002; Meier et al. 2006). The development of DNA-based approaches shows promise for overcoming these problems, and many taxonomists have adopted an integrative taxonomic approach (Dayrat, 2005), which allows combining data from multiple lines of evidence such as morphology, ecology and genetics. While an integrative taxonomic approach delivers a much-needed rigor in taxonomic practices, the disagreements still remain as to what kind of data should be prioritized and the degree of congruence to be expected from these different datasets (Padial et al. 2010; Schlick-Steiner et al. 2010; Yeates et al. 2011). Therefore, there still remains the need for a more objective framework and quantifiable criteria for defining species boundaries (Pante et al. 2015).

Geographically widespread taxa pose a particular challenge for species delimitation as their distributions often tend to be uneven and consist of populations with allopatric ranges (Mutanen et al. 2012). In cases of allopatric populations, it is difficult to assess the reproductive compatibility and the decision to define genetically separated populations as distinct species depends greatly on the species concept applied and taxonomists’ subjective ideas of species boundaries and taxonomically important features (Funk and Omland 2003), making species delimitation of allopatric populations arbitrary and often unstable. Species distributed in mountainous habitats form a classic example of patchy distributions with frequent allopatry. Arctic-alpine species provide an excellent model to study the allopatry problem as their disjunct distributions are typically the result of repeated cycles of range expansion and contraction driven by the glacial oscillations during the Quaternary (Hewitt 2004). These cold-adapted species evidently had a widespread, contiguous distribution during ice-ages and as a result of increased postglacial warming, shifted towards north into the boreal, subarctic and arctic belts and into mountain areas in more southern parts, resulting in an increased spatial separation (Schmitt 2007). For Holarctic taxa, the distributions are particularly large and often encompass the entire northern hemisphere (Schmitt et al. 2010). There are plenty of studies that have examined patterns and levels of genetic divergence between taxa distributed in arctic and alpine biogeographic zones (Muster and Berendonk 2006; Schmitt et al. 2006, 2016; Schmitt 2007; Hou et al. 2015; Lindholm et al. 2016; György et al. 2018). However, these studies usually have focused on small geographic regions and have almost always been based on limited numbers of genetic markers.

Accumulation of massive barcoding libraries along with a comparatively well-established taxonomy has allowed taxonomists to study patterns of genetic divergences of the standard cytochrome oxidase I (COI) gene of mitochondrial DNA (mtDNA) at large geographic scales for megadiverse insect orders such as Coleoptera and Lepidoptera (Bergsten et al. 2012; Huemer et al. 2014, 2018). These studies have shown that in mtDNA, intraspecific divergences tend to increase with sampling intensity and geographic distance. Cases of deep intraspecific barcode divergences in arctic and alpine taxa, as well as between Alps/Fennoscandia vs. North America might point to potentially overlooked species (Mutanen et al., 2012). Several other studies also revealed inconsistencies in so-called ‘barcode gaps’ in well-sampled groups (Meyer and Paulay 2005; Wiemers and Fiedler 2007), highlighting the need to examine the thresholds of intra- and interspecific variation in more detail using nuclear loci. While curated DNA barcode libraries are useful in rapid species discovery and are also known to be effective in revealing potential cryptic species (Mutanen et al. 2016), such a single-locus approach suffers from limitations, e.g., because of retention of ancestral polymorphisms, male-biased gene flow, asymmetrical introgression, paralogy and *Wolbachia*-mediated genetic sweeps (Funk and Omland 2003; Ballard and Whitlock 2004; Moritz and Cicero 2004), rendering the COI gene trees to deviate from those of the species.

As the cost of sequencing continuously decreases, genomic-scale data generated through high-throughput sequencing technologies is now increasingly used in both phylogenetic inference and species delimitation (Herrera and Shank 2016; Breinholt et al. 2018; Grummer et al. 2018; Pie et al. 2019). There is also a suite of methods, developed in statistically rigorous frameworks such as multispecies coalescent and Bayesian frameworks (e.g., Leaché et al., 2014; Yang, 2015), that show promise for bringing more objectivity to species delimitation. Additionally, simulations have shown the increasing accuracy and robustness of these methods in the presence of different biological processes such as gene flow and incomplete lineage sorting etc. (Jackson et al. 2017; Zhang et al. 2018). There are also newer sets of software that are faster and scalable to larger datasets (Fujisawa et al. 2016; Rabiee and Mirarab 2020), which can overcome the computational limitations imposed by Bayesian programs. Sequence capture methods such as target enrichment can be useful in bringing stability and even common standards for species delimitation if based on a fixed set of loci that are found across a large diversity of taxa. This idea was recently put forth by Eberle et al. (2020) and tested in a selection of metazoan taxa by Dietz et al. (2022).

We here use genomic scale data to study intraspecific divergence patterns on a broad geographic scale. More specifically, we use a target enrichment approach to understand the intraspecific patterns and to delimit the allopatric populations of ten species of Lepidoptera with an arctic-alpine and Holarctic distribution. These species were deliberately selected to represent various geographic patterns in their COI gene across their geographic range. We first examined if COI divergences provide a good proxy for the genome-wide divergences by comparing uncorrected pairwise distances and phylogenetic structure between the allopatric populations of the focal species. We secondly investigated the efficiency of the target enrichment approach in delimitation of allopatric populations. To reach this goal, we made use of population genetic, multispecies coalescent and recently developed species delimitation approaches designed for genomic data. Our sampling includes individuals from four major regions across the Holarctic realm – Fennoscandia (Northern Europe), Alps (with a few samples from Apennine mountains and Pyrenees, Central and southern Europe), Altai mountains (Asia) and the Nearctic (North America). We divided our species groups into two categories. In the first category, we focused on comparing genetic patterns across allopatric populations within the Palearctic region (intra-continental). In the second category (intercontinental), we also included populations from North America (the Nearctic region).

With our study, we aim to provide insights on genetic patterns for a set of widely distributed Palearctic and Holarctic taxa, as well as initiate discussion for determining and establishing better justified and repeatable standards for delimitation of allopatric populations by genomic means, as we find their delimitation presently lacking a solid epistemological foundation.

## MATERIALS AND METHODS

### Study species and sample preparation

Species for the intra-continental category include *Eudonia sudetica* (Zeller, 1839) (Crambidae) with a disjunct arctic-alpine distribution pattern in Europe, and *Elophos vittaria* (Thunberg, 1788) (Geometridae), *Syngrapha hochenwarthi* (Hochenwarth, 1785) (Noctuidae) and *Xestia speciosa* (Hübner, 1813) (Noctuidae), with patchy Palearctic distribution patterns. The species included in the intercontinental category include *Agriades glandon* (de Prunner, 1798) (Lycaenidae)*, Arctia caja* (Linnaeus, 1758) (Erebidae)*, Carsia sororiata* (Hübner, 1813) (Geometridae)*, Coenonympha tullia* (Müller, 1764) (Nymphalidae)*, Macaria brunneata* (Thunberg, 1784) (Geometridae), and *Xestia lorezi* (Staudinger, 1891) (Noctuidae).

Samples were collected from various sites across four main regions (Fig. 1). Table 1 gives the number of taxa and an overview of the populations analyzed for each species. Further details regarding the sample localities (wherever the information was available) is given in the supplementary file (Table S1). Genomic DNA was extracted from either the thorax or legs of dry specimens. DNA extraction was performed using the QIAGEN DNeasy Blood and Tissue Kit (Hildesheim, Germany) or the Omega Bio-tek E.Z.N.A. Insect DNA Kit (Norcross, United States) following the protocols given by the manufacturers.

**Figure 1:**
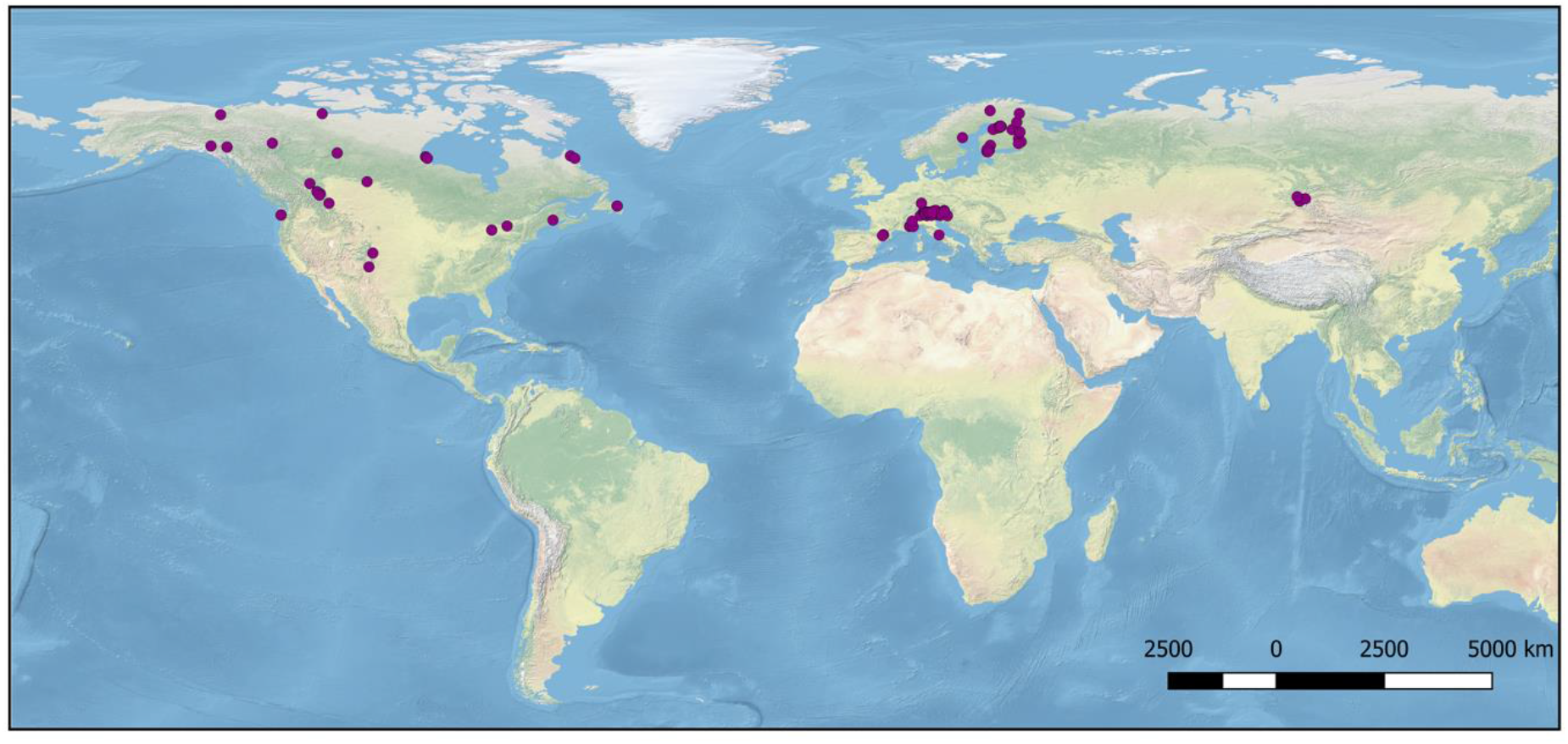
The map of sampling localities

**Table 1:**
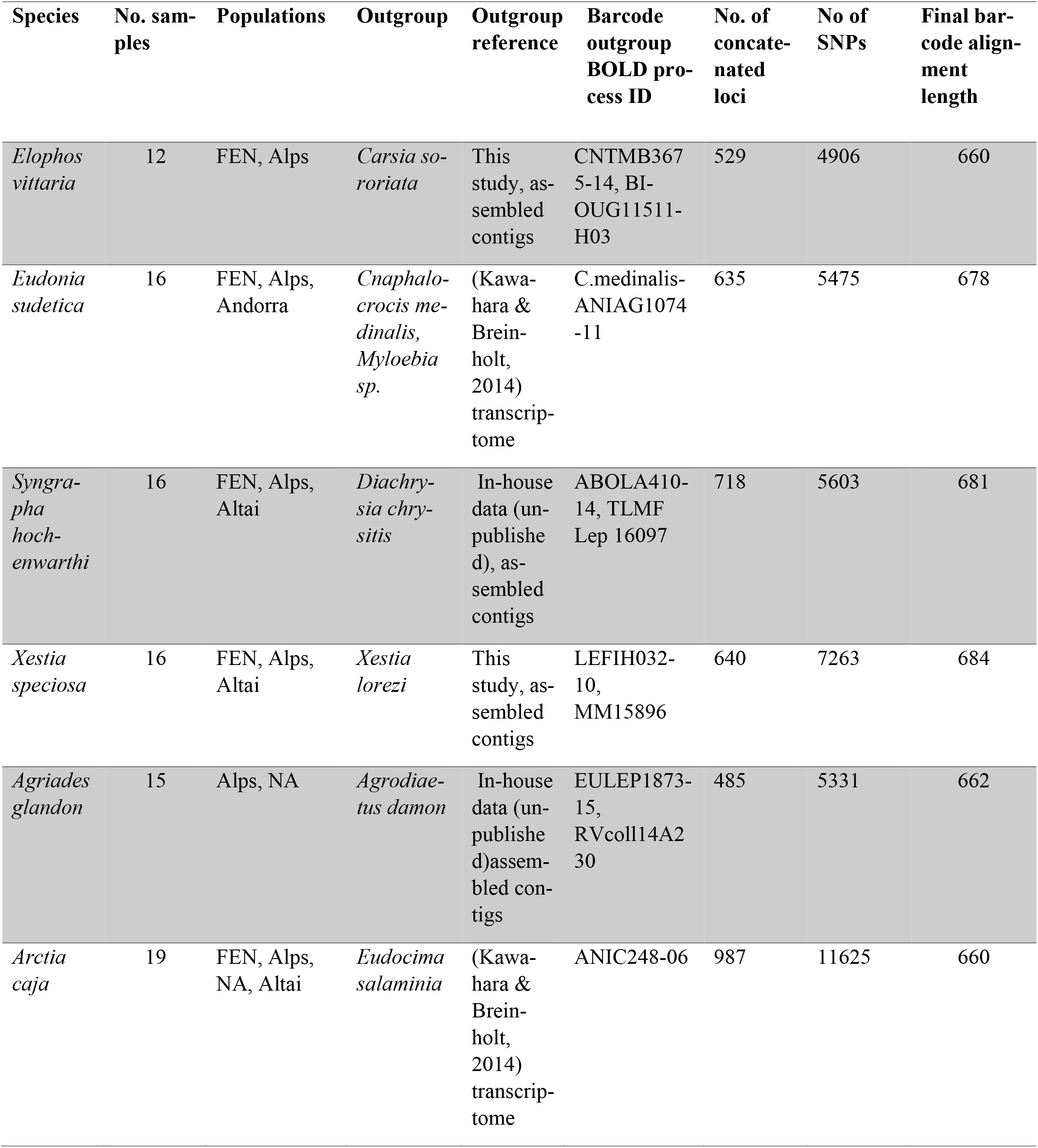
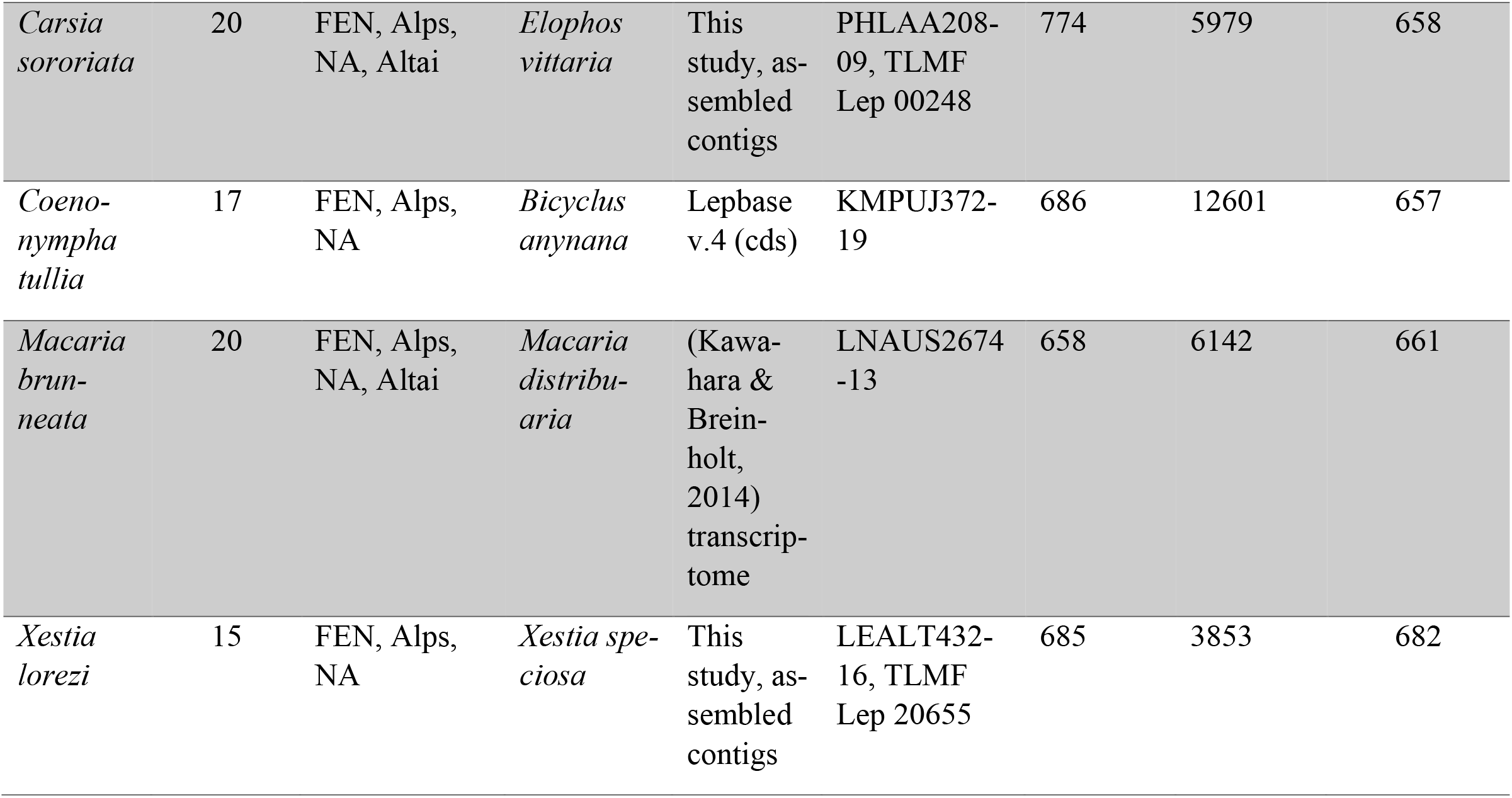
No. of samples included for each species group, populations sampled, outgroup information, No. of SNPs and the final barcode alignment length

### Target enrichment laboratory procedures

Following extraction, the DNA concentration of each sample was measured using the Invitrogen Quant-iT PicoGreen dsDNA (Waltham, United States) quantification kit. Approximately 100 ng of absolute DNA was taken for further processing as per the standard Agilent protocol; for the samples with low DNA yield, approximately 50 ng or 10 ng of absolute DNA was used, depending on the available quantity. The genomic DNA was subjected to random mechanical shearing to an average size between 150-200 base pairs (bp) using the Diagennode Bioruptor sonication device. This was followed by end repair reaction and ligation of adenine residue to the 3’ end of the blunt fragments (A-tailing) to allow ligation of barcoded adaptors using the Agilent SureSelect XT HS2 DNA library preparation kit (Santa Clara, United States). PCR amplification of adaptor-ligated libraries was then performed, after indexing with the Agilent SureSelect XT HS2 primer pairs; we used 8 PCR cycles for all of the samples, irrespective of the amount of absolute DNA taken initially.

After the library construction, custom Agilent SureSelect baits (6-11.9 Mb) were used for solution-based target enrichment of a pool containing 8 or 16 libraries. The hybridisation was performed on each pool with the Agilent SureSelect XT HS2 Target Enrichment Kit ILM module following the manufacturer’s instructions. Enriched libraries were then captured with MyOne Streptavidin T1 beads. These captured libraries were PCR amplified and the final concentration of each captured library was measured using the Invitrogen Quant-iT PicoGreen dsDNA quantification kit. Following the enrichment, pooled libraries were sequenced using the Illumina Nextseq 500 platform (mid output) to generate paired-end 150- bp reads at FIMM Genomics (Institute for Molecular Medicine Finland), Helsinki.

### Bioinformatics processing

The raw demultiplexed data received from the sequencing provider were processed with the TEnriAn (Target ENrichment ANalysis) pipeline (Mayer et al. 2021). Briefly, this pipeline has three main steps of data analysis. In the first assembly step, raw reads are merged, trimmed to remove adaptor and low quality bases using fastp (Chen et al. 2018) and are assembled with TRINITY (Grabherr et al. 2011). After the assembly, samples are checked and filtered for potential cross-contaminations using ‘all against all’ blast searches. However, we avoided doing this step in our analysis as we observed that it was removing a lot of useful data, since our samples are very closely related, and high similarity between samples is expected. In the second step, specimens are assigned to orthologous clusters. For this step, the Orthograph (Petersen et al. 2017) sets were prepared using official gene sets (OGS) of five species (*Bombyx mori*, *Danaus plexippus*, *Heliconius melpomene*, *Melitaea cinxia*, *Papilio glaucus*). After the sets were generated, the orthology step was run to generate clusters of orthologous genes. In the third step, these clusters inferred for each specimen are aligned against reference Hidden Markov Models (HMMs) and are filtered using various criteria such as coverage, Aliscore (Misof and Misof 2009), outliers, etc. The final alignments produced were manually checked in Geneious Prime (https://www.geneious.com/prime/) to make sure that there were no misalignments. We further removed alignments with a GC content higher than 60% before proceeding ahead, as those can have a negative effect on downstream phylogenetic analyses.

### Barcoding

We analyzed mitochondrial DNA barcodes for selected samples for each of the ten case studies. Several barcodes were publicly available for the ten species and were retrieved from BOLD. For reminder of the samples, we sequenced the mitochondrial COI region from DNA extracts in the lab, the protocol details of which can be found in the supplementary file. For the samples that barcoding was unsuccessful on, we extracted the barcode region from the target enrichment data using *exonerate* ver. 2.4.0 (Slater and Birney 2005) using the alignment of publicly available and new barcodes as a query, and our de-novo assembled target enrichment assemblies as a target. Details of the commands are given in the supplementary file. These extracted barcodes were then aligned with the rest of the barcodes for each study species in Geneious Prime using Muscle ver. 5.1 (Edgar 2004). The final barcode alignment lengths for each species can be found in Table 1. All the newly generated barcodes were deposited on BOLD after processing.

Phylogenetic trees of the aligned COI sequences were generated using IQTREE ver. 2.0.3 (Minh et al. 2020), using the best-fit model estimated by the program using ModelFinder (Kalyaanamoorthy et al. 2017) and ultrafast bootstrapping with 1000 replicates. We used -bnni option to reduce the impact of model violation and overestimated branch support. The phylogenetic analysis was run 20 times for each of the study species to assess the congruence among the tree searches. For the sake of consistency, for each species we included the same outgroup that was chosen for rooting the target enrichment phylogenetic trees. The details of the outgroup selection are given in Table 1. The trees with the highest likelihood were visualized in Figtree v1.4.4 (available from <u>tree.bio.ed.ac.uk/software/figtree/</u>) and rooted on the selected outgroup.

### Phylogenetic analyses

For target enrichment data, we carried out phylogenetic analyses using both concatenation and coalescent approaches. Maximum Likelihood (ML) phylogenetic analyses for the target enrichment data were performed using IQTREE ver. 2.0.3 (Minh et al. 2020). In order to partition the data according to gene positions, we set up partitioned analysis using IQTREE with option -m TESTMERGEONLY to resemble PartitionFinder (Chernomor et al. 2016) and the rcluster algorithm (Lanfear et al. 2014) with the rcluster percentage set to 10, under the AICc criterion. The best partitioning scheme was then used as an input to set up a partitioned analysis in IQ-TREE. We used the ultrafast bootstrap approximation with 1000 replicates (Hoang et al. 2018). We also performed a SH-like approximate likelihood ratio test (Guindon et al. 2010) with 1000 bootstrap replicates using the -alrt option. To further reduce the risk of overestimating branch supports, the -bnni option was used. Each analysis was run 20 times to assess the congruence among the tree searches. The tree with the highest likelihood was visualised in Figtree and phylogenetic trees were rooted on their selected outgroups (Table 1)

In addition to ML analyses, we did species tree analyses in ASTRAL-III v. 5.7.4 (Zhang et al. 2018), which is statistically consistent under the multispecies coalescent framework and takes into account incomplete lineage sorting (ILS) while inferring the species tree. Model selection on each individual gene alignment for each of the species was performed using ModelFinder (Kalyaanamoorthy et al. 2017) and tree inference was done in IQTREE. The gene trees were further filtered using program TreeShrink (Mai and Mirarab 2018) before running ASTRAL to detect and prune the abnormally long branches. the program was run in default ‘per-species’ mode with a false positive tolerance rate (α) set to 0.05. The resulting shrunk output gene trees were used as an input for ASTRAL, which generated a species tree along with the quartet score.

### Uncorrected P-distances

We calculated the uncorrected P-distances using PAUP* ver. 4 (Cummings 2004) for target enrichment concatenated dataset and the barcodes, excluding the outgroup from the alignment. The goal was to determine the percent pairwise genetic distances between populations of the same species using both barcodes and target enrichment, and to compare these distances to determine the level of agreement. The calculated distances were grouped into different categories which included comparing individuals from each population with one another as well as within population comparisons. The absolute values of distances were visualized as boxplots using the R package *ggplot2* (Wickham 2016).

### SNP calling

For each of the ten study species, we used the ingroup samples with the highest number of orthologous clusters inferred (during orthology assessment step in the TEnriAn workflow) as a reference for SNP calling, similar to the procedure mentioned by Zarza et al. (2016) and Erickson et al. (2020) for UCE data. The raw, cleansed data (i.e., after removing adaptors and low-quality bases) were mapped against this reference using with the BWA-MEM algorithm in bwa 0.7.17 (available from <u>bio-bwa.sourceforge.net</u>) with the minimum seed length set to 30.

SNPs were then extracted from the sorted BAM files generated during mapping by using the samtools *mpileup* option piped together with the bcftools *call* option. Filtering was then completed using bcftools, in the first step to keep only biallelic SNPs filtering out indels and multiallelic SNPs, and in the second step to keep one randomly chosen SNP per locus. The final number of SNPs retained for each species after the first filtering step is mentioned in Table 1.

### Population genetics analyses

For each study species, we performed a principal component analysis (PCA) on the SNP dataset using the *dudi.pca* function from the R package *adegenet* (Jombart and Ahmed 2011). Additionally, we calculated pairwise F_ST_ for each population pair in each of the ten study species using R package *hierfstat* (Goudet 2005). F_ST_, which is a measure of population differentiation, quantifies the differences in allele frequencies among the populations.

We also performed a STRUCTURE analysis (Pritchard et al. 2000) for species with more than two populations (except for *X. lorezi*, as we did not have sufficient sampling from North America) to infer the presence of admixture. For this, our target enrichment SNP data in vcf format was converted to structure format (.str) using PGDSpider ver.2.1.1.5 (Lischer and Excoffier 2012). We tested five putative numbers of clusters, K=1–5, with 10 iterations for each K. To determine the optimal number of genetic clusters (K), we used the ΔK method in STRUCTURE HARVESTER (Evanno et al. 2005; Earl and vonHoldt 2012) with 500 000 generations for the Markov chain and a value of 100 000 as burn-in. These cluster assignments were then aligned across all 10 replicates and visualised using CLUMPAK (Kopelman et al. 2015) server. The optimum K inferred for each species and their respective ΔK plots are given in the supplementary file.

### Isolation by Distance (IBD)

We tested for isolation by distance (IBD) using a Mantel test between a matrix of genetic distances and matrix of geographic distances using *mantel.randtest* function in R package *adegenet*. This test finds the correlation between individual Edwards’ genetic distances and Euclidean geographic distances (Mantel 1967). Based on the simulated p-value, we determined whether isolation by distance was significant (p < 0.05). This analysis was mainly carried out for Holarctic taxa *X. speciosa*, *A. caja*, *C. sororiata*, *C. tullia* and *M. brunneata*. The histograms of simulated p-values and the isolation by distance scatterplots are given in the supplementary file.

### Species delimitation

To test for the presence of species level differentiation in our datasets, we performed species delimitation analyses using two different programs. To test the alternate scenarios where different populations consist of a single or distinct species, we used program *tr2,* a multilocus species delimitation method that finds the best delimitation based on a distribution model of rooted triplets (Fujisawa et al. 2016). In each of the ten cases, we used gene trees rooted using the newick utilities package (Junier and Zdobnov 2010) as an input. Each scenario was compared against the null model, which assumes the presence of single species, based on -log(likelihood) scores, and the model with lowest score was inferred as the best delimitation model, hence the best delimitation scenario. The details of different models corresponding to different scenarios tested for each species group and their -log(likelihood) scores are given in the supplementary file.

We also tested a recently developed quartet-based species delimitation method SODA (Rabiee and Mirarab 2020), which is designed to be scalable to large datasets. For this, we used the same set of rooted trees as *tr2*. We ran two different analyses using SODA, first using the tree generated from ASTRAL as a guide tree and without mapping individuals to populations. The second analysis was run using a mapping file with population groupings specified a priori. The output of this program consists of each individual assigned to the distinct species.

## RESULTS

### Dataset overview

The total number of concatenated loci after various filtering steps for each species can be found in Table 1. The number of informative loci and the missing data for each individual from all the species can be found in Table S1. The highest number of informative loci were captured for *A. caja* (avg. no. of loci = 868, std. dev. = 35.8) and lowest for *A. glandon* (avg. no. of loci = 406, std. dev. = 18.3). The highest percentage of missing data was observed in *C. tullia* (avg. = 33.9%, std. dev. = 0.11, Table S1) and lowest was observed in *A. caja* (avg. = 10.6%, std. dev. = 0.06, Table S1).

### P-distance comparisons

Figure 2a shows the P-distance boxplot comparisons between barcode and target enrichment for species studied under intra-continental category. For *E. vittaria*, the median values of both barcodes and target enrichment P-distances within Alps and Fennoscandian populations fell between 0-1.5%. In this species, however, the comparison varied greatly for genetic distances between Alps and Fennoscandia. While the median value for target enrichment distances fell between 0.15-2%, for barcodes it was more than 5%. For *E. sudetica*, we observed median values of barcode P-distances for Alps/Apennines-FEN and Alps/Apennines-Pyrenees comparison close to 2%, but remarkable variation in barcode P-distance values for within Alps/Apennines and no variation for within-FEN category. This large variation, as well as the outliers were barcode P-distances reaching near 6% and caused by a divergent individual from Italy/Apennines (TLMF Lep 03039; note this individual clusters with four barcoded specimens from central Italy). Target enrichment P-distance values on the other hand remained below 2%, with median P-distance values for all categories near 1% and for within FEN comparison less than 1%. For *S. hochenwarthi*, medians for both distances across all the categories were below 1%, but the differences in the medians of both barcode and target enrichment distances for all the categories were significant. In general, the target enrichment distances tended to fall between 0.5-1%, which was not the case for barcode distances, except for Alps-Altai and FEN-Altai category. For *X. speciosa* which is known to have a barcode split in Europe, we observed barcode distances greater than 2% for FEN-Alps and Alps-Altai category (except for outliers observed for the first category) and between 0.15-2.5% FEN-Altai category. To the contrary, target enrichment P-distance values tended to remain below 1% for all the categories.

**Figure 2:**
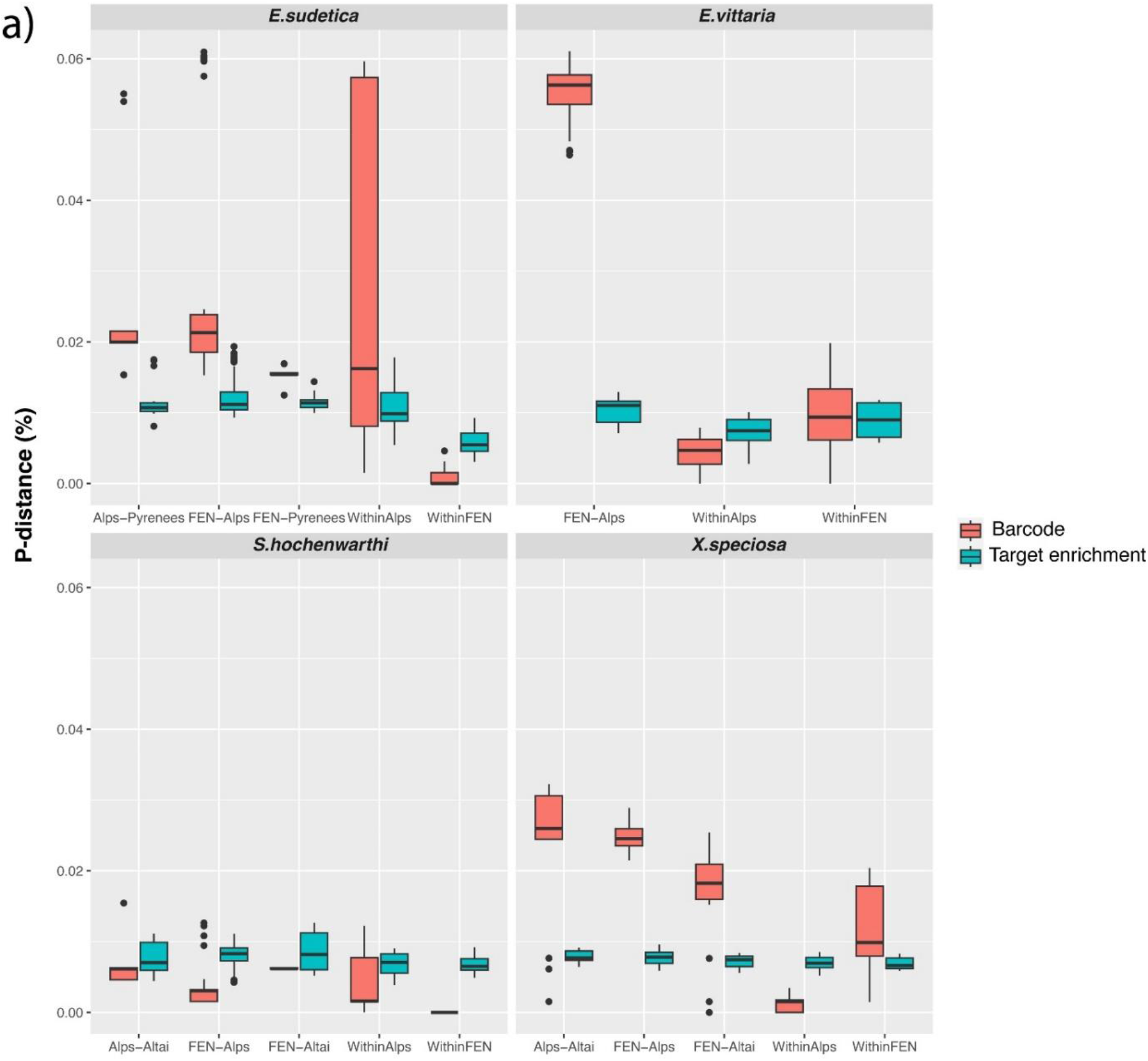
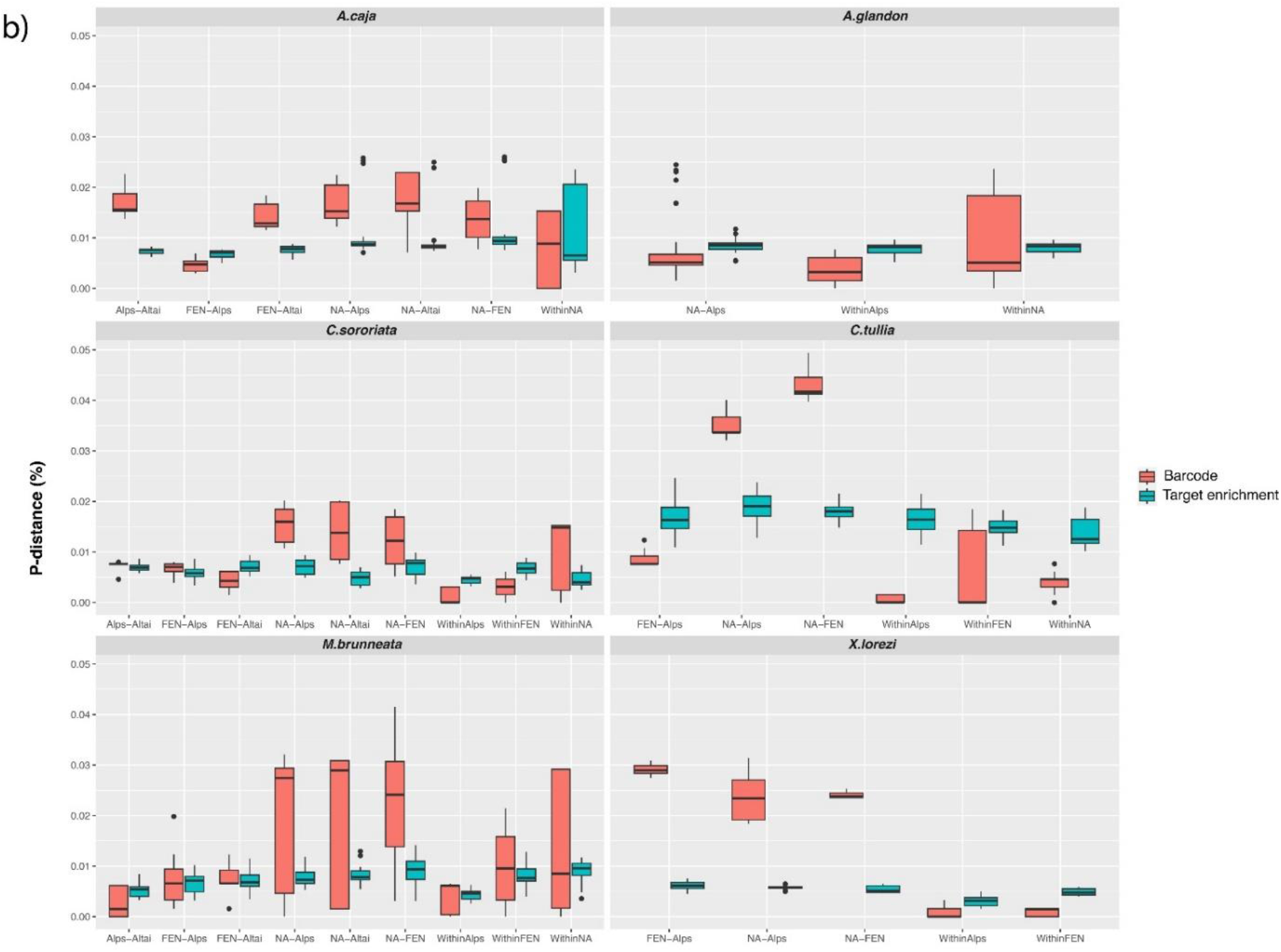
P-distance comparison boxplots for a) Species included in intra-continental category b) Species included in intercontinental category

For the taxa in the intercontinental category (Fig. 2b), we observed a larger variation in barcode P-distance values than the target enrichment P-distances. This is observed for *A. glandon*, *A. caja*, *C. sororiata* and *M. brunneata*. We also observed that the barcode P-distance values for the within NA category regularly varied greatly. The only exception for this pattern was *C. tullia*. The target enrichment P-distance values for Holarctic taxa tended to stay below 1%, again except for *C. tullia*, where they were all above 1%. In general, we observed that when the North American population (referred to as NA population henceforth) was included, the barcode P-distance values tended to have a greater variation. For *X. lorezi*, all the three populations – NA, Alps, and Altai showed deep barcode splits (more than 2%). A similar deep split was observed in barcodes for *C. tullia*, where distance between Nearctic and both Palearctic populations was greater than 3%. Similarly for *M. brunneata*, the median barcode distances of Nearctic population from each of the Palearctic populations were more than 2%. We did not observe deep splits in barcodes in *C. sororiata* and *A. caja*. For *A. glandon*, median barcode P-distances were less than median target enrichment P-distances.

### Phylogenetics and population genetics – Intra-continental scale

*Elophos vittaria*: This species shows a typical disjunct arctic-alpine distribution pattern. The ASTRAL tree supported the divergence between the Alps and Fennoscandia (quartet support value of 1 for Fennoscandian population and 0.95 for the Alps population, Fig. 4a), which was also observed in previous studies. However, barcode and ML trees did not match this pattern. In barcode tree, we observed the monophyly of Alps population but not Fennoscandian population and in ML tree only Fennoscandian population was observed to be monophyletic (Fig. S1a). In the barcode tree, the individual MM06327 clusters together with the Alps population, suggesting that the barcode sequence extracted from the target enrichment data could be chimeric. This individual also shows somewhat longer branch length in the ML tree and shows a high percent of missing data as compared to the other individuals (Table S1). In the PCA, individuals from the Alps were clustered together more closely while the Fennoscandian individuals were observed to be far apart (Fig. 3a). The pairwise F_ST_ between two populations was observed to be 0.279 (Table 2).

**Figure 3:**
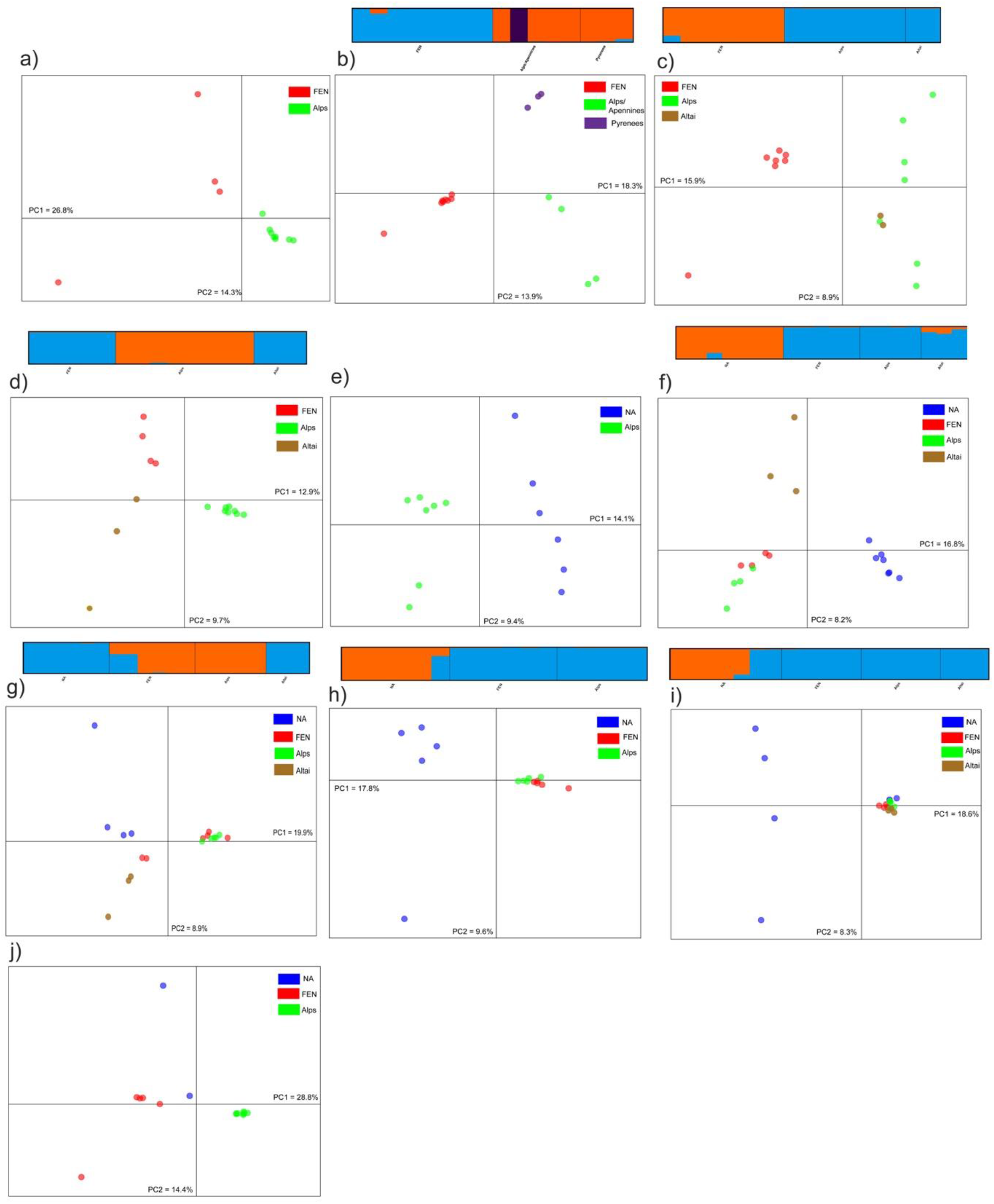
PCA of a) *E. vittaria* e) *A. glandon* j) *X. lorezi*. PCA and STRUCTURE K=2 or K=3 barplot for b) *E. sudetica* c) *S. hochnewarthi* d) *X. speciosa* f) *A. caja* g) *C. sororiata* h) *C. tullia* i) *M. brunneata*

**Figure 4:**
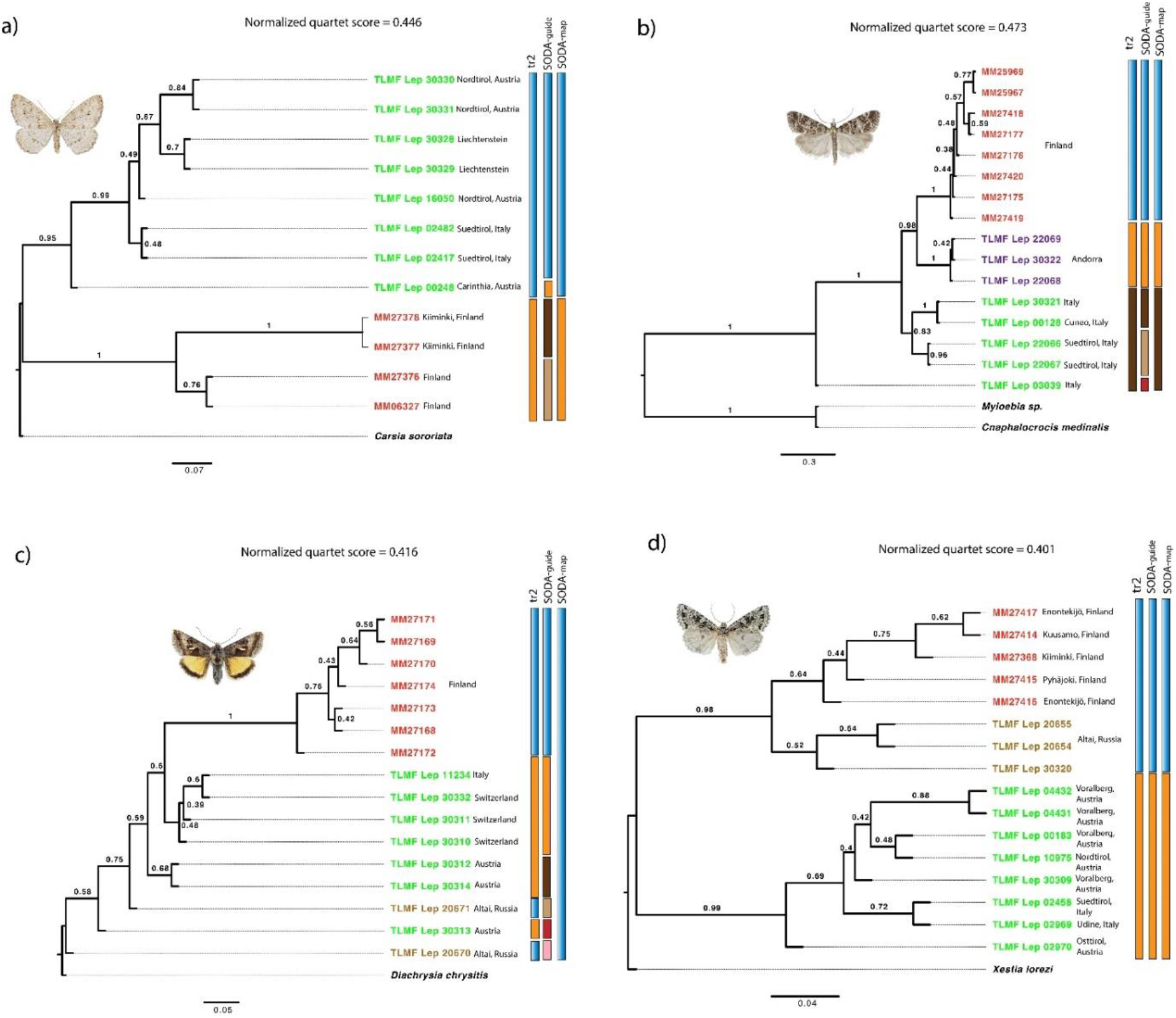
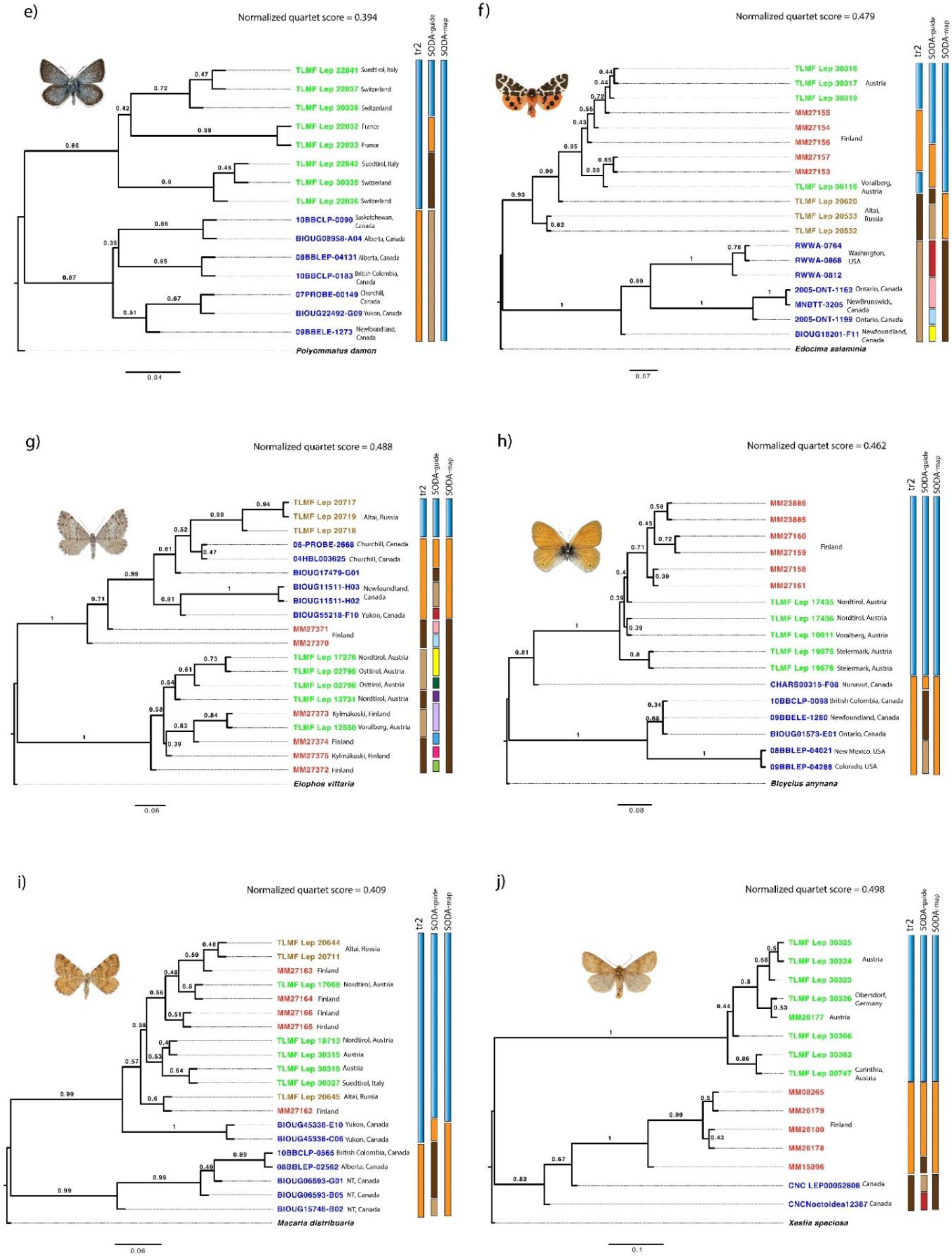
Species delimitation analyses mapped onto ASTRAL tree for a) E. vittaria b) E. sudetica c) S. hochenwarthi d) X. speciosa e) A. glandon f) A. caja g) C. sororiata h) C. tullia i) M. brunneata j) X. lorezi

**Table 2:**
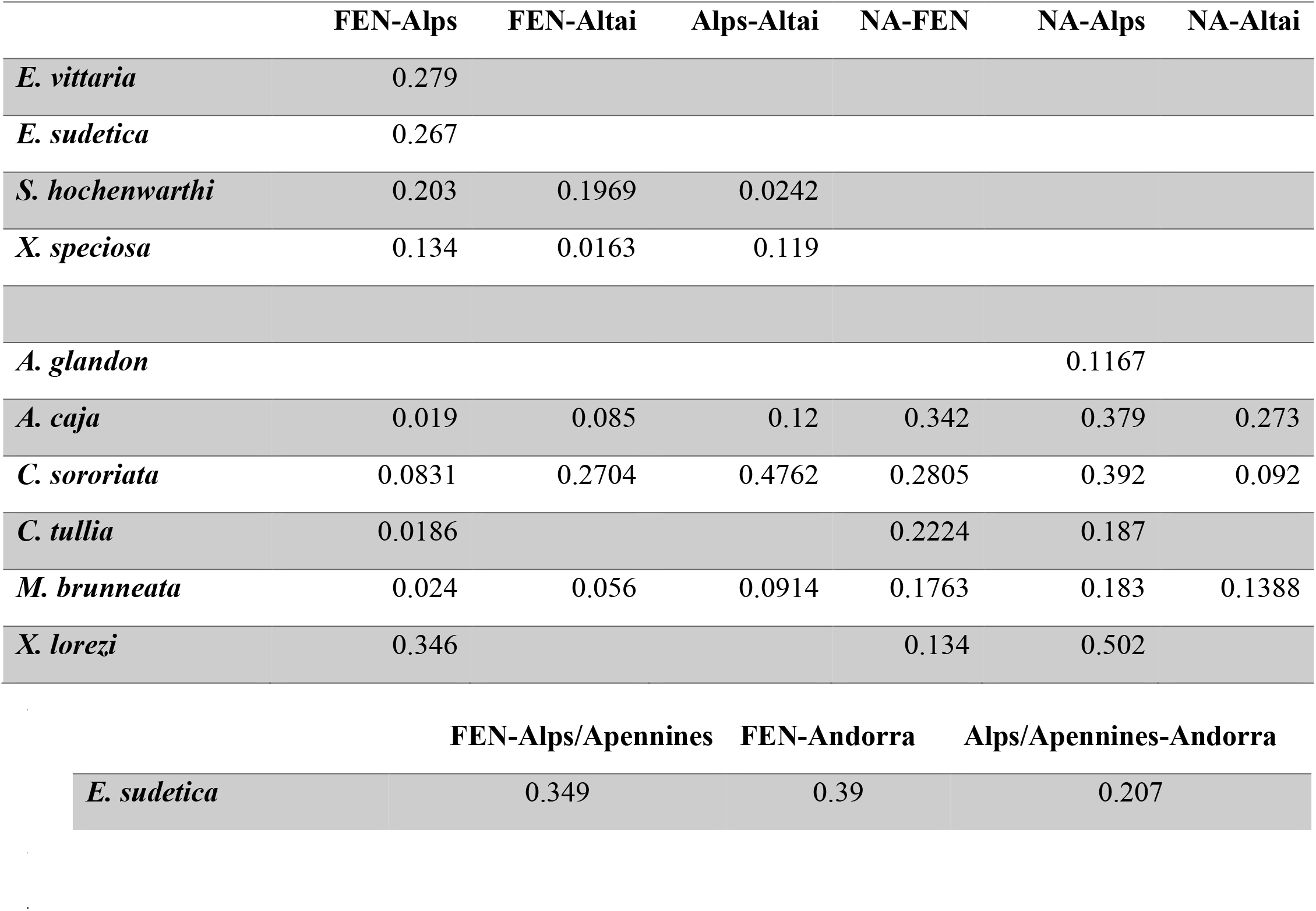
Pairwise F_ST_ values for each population comparison

*Eudonia sudetica*: The phylogenetic relationships between *E. sudetica* populations were observed to be varying between the three approaches, although in all three cases, Fennoscandian specimens clustered together (with support value of 89% in barcode tree, 100/100 in ML tree and 0.99 in ASTRAL, Fig. 4b and S1b). Additionally, in all the three cases, the individuals sampled from the Pyrenees also clustered together with a high support value in ML and ASTRAL trees (Fig. 4b and S1b, 99.5/99 in ML and 1 in ASTRAL). However, the phylogenetic placement of this and the Alps/Apennines population differed in all the three trees. While in the barcode tree the Alps/Apennines population is inferred as a sister group to the Fennoscandian population, in the ML tree the individuals from this population are paraphyletic with respect to the clade of Fennoscandian and Pyrenees individuals. In the ASTRAL tree, this population forms a sister clade to the other two. In the barcode tree, the clade of Pyrenees individuals is shown to be sister to the cluster of both Fennoscandian and Alps individuals (except for Apennines individual), while in ASTRAL it shown to be sister to a cluster of Fennoscandian individuals. All the trees showed the individual TLMF Lep 03039 from Apennines as divergent, forming a sister to all the rest of the individuals. In PCA, separation of Fennoscandian and the southern populations along PC1 is observed, and so is the separation of Pyrenees individuals along PC2 (Fig. 3b). STRUCTURE results revealed K=3 as optimum number of clusters for this dataset, with Fennoscandian population as one genetic cluster, Alps and Pyrenees population as another and the Apennines individual as a third separate genetic entity (Fig. 3b). Relatively high pairwise F_ST_ between Fennoscandian and each southern population was also present (Table 2).

*Syngrapha hochenwarthi*: This species is characterized by having a disjunct Palearctic distribution. Our results supported the Fennoscandian population as a separate cluster in all the barcode, ASTRAL and ML trees (support values of 78%, 1, 100/82 respectively, Fig. 4c and S1c). The remainder of the individuals, belonging to the Alps and Altai populations, were observed to be paraphyletic with respect to the Fennoscandian cluster. PCA and STRUCTURE also supported genetic distinctiveness of Fennoscandian population (Fig. 3c). The pairwise F_ST_ between Fennoscandia-Altai and Fennoscandia-Alps was observed to be around 0.2 (Table 2).

*Xestia speciosa*: We observed that the specimens from the Alps cluster together in barcode and ASTRAL trees, with support values of 95% and 0.99 respectively, whereas the ML tree shows a mixed pattern (Fig. S1d). Even though ASTRAL clusters specimens from Fennoscandia and Altai population with quartet support value of 0.98, the support values for each population as a distinct lineage are low (Fig. 4d). The PCA and STRUCTURE further supports the genetic distinctiveness of the Alps population (Fig. 3d), and the pairwise F_ST_ values compared to the Fennoscandian and Altai populations are 0.13 and 0.12, respectively (Table 2). As expected, Fennoscandian and Altai population show low pairwise F_ST_ of 0.06. Testing for Isolation by distance using a Mantel test was not significant (p-value > 0.05).

### Phylogenetics and population genetics – Intercontinental scale

*Agriades glandon*: We observed discordance between the phylogenetic trees obtained from barcodes and the genomic data. While barcodes did not show genetic separation of the Alps and NA population, ASTRAL inferred them as two separate clusters with quartet support value of 0.86 for the Alps and 0.87 for NA population (Fig. 4e and S1e). We also observed that tree topology of target enrichment ML tree was largely different from that of ASTRAL. In the PCA, the two populations showed separation mainly along PC1 (Fig. 3e) with the pairwise F_ST_ between two populations calculated as 0.12 (Table 2).

*Arctia caja*: We observed discordance between phylogenetic relationships inferred from barcodes and the genomic data. Phylogenetic trees inferred from both ML and ASTRAL approaches using genomic data differed only slightly, i.e., mainly in the relationships of populations within Eurasia. Both the ML and ASTRAL trees supported the separation of the NA cluster with high support (99/100 in ML and quartet support value 1 in ASTRAL, Fig. 4f and S1f). In the PCA, we observed that the separation of NA from the Eurasian samples was mainly explained along the PC1, whereas the separation of Altai samples from the rest of the populations was observed along PC2 (Fig. 3f). We also observed high pairwise F_ST_ values between NA and the remainder of the populations, especially for Fennoscandian and Alps populations it was above 0.3 (Table 2). For the NA and Altai populations, the pairwise F_ST_ value was 0.27. Signs of admixture are present, possibly caused by the historical contact, mainly between NA and Altai population as shown by the STRUCTURE results (Fig. 3f). The Mantel test revealed significant isolation by distance (p-value < 0.05).

*Carsia sororiata*: For this species, which has a Holarctic distribution with boreo-montane disjunction, we did not observe any clear geographic pattern in any of the approaches. While samples from Altai clustered together in ML and ASTRAL trees (target enrichment ML tree with support values of 100/100 and in ASTRAL with quartet support value of 0.99, Fig. S1g and 4g), no such observation was made for other populations. In barcode tree, only individuals from the Alps clustered together (support value 84%, Fig. S1g), while those from other three populations did not form clear groups. Both the ASTRAL and target enrichment ML trees supported the close phylogenetic relatedness of the Altai and North American populations. The same can be observed from the STRUCTURE results, where NA and Altai population belong to the same genetic cluster (shown in orange in Fig. 3g). While the Alps and Fennoscandian populations form second genetic cluster (shown in blue in Fig. 3g). Two of the Fennoscandian individuals show signs of admixture with the NA-Altai genetic cluster. Their placements are also different in the ASTRAL and ML trees (individuals MM27371 and MM27370), where they were part of the NA-Altai clade. In the PCA, these two samples were rather far away from the cluster of the remainder of the Fennoscandian and Alps individuals (Fig. 3g). We observed pairwise F_ST_ of 0.28 and 0.39 between Fennoscandian-NA and Alps- NA populations, respectively. F_ST_ values of 0.27 and 0.48 were observed for Altai- Fennoscandia and Altai-Alps, respectively. We observed relatively low pairwise F_ST_ between NA and Altai populations (0.092, Table 2). Isolation by distance was observed to be significant (p-value < 0.05).

*Coenonympha tullia*: Barcode and ASTRAL trees revealed the presence of two separate clusters corresponding to Fennoscandian and NA populations (Fig. S1h and 4h), and the Alps individuals as paraphyletic with respect to Fennoscandian population. However, the target enrichment ML tree did not show this pattern where only Nearctic population formed a separate genetic cluster (support value 100/100) except for an individual from Nunavut, Canada (CHARS00315-F08) and all the reminder of the individuals were observed to paraphyletic with respect to this NA clade (Fig. S1h). The STRUCTURE results support the genetic distinctiveness of the NA population, and the Nunavut individual seems to be admixed with the gene pools from both Europe and North America (Fig. 3h). Whether this is the true admixture or the result of cross-contamination during lab work remains unclear.

Pairwise F_ST_ values between NA and each of the Fennoscandian and Alps populations were 0.22 and 0.19, respectively. The clustering of Fennoscandian and Alps individuals in the PCA (shown in red and green in Fig. 3h), as well as the low F_ST_ observed between the two indicates no genetic separation. The Mantel test revealed that the isolation by distance is not significant (p-value > 0.05).

*Macaria brunneata*: For this species, we observed that all the three approaches supported separation of the NA population, with support values 82% in the barcode tree, 0.99 in ASTRAL and 85.9/84 in the ML tree (Fig. 4i and S1i). However, two individuals from Yukon, Canada did not cluster together with the NA clade, but were rather part of the Eurasian clade consisting of individuals from the Alps, Fennoscandia and Altai. These individuals seem to have a Eurasian gene pool as suggested by STRUCTURE and PCA (Fig. 3i). The pairwise F_ST_ between the NA and Fennoscandian populations as well as between the NA and the Alps populations were 0.18, and that between the NA population and Altai was 0.14. The Mantel test revealed significant isolation by distance (p-value < 0.05).

*Xestia lorezi*: In case of arctic-alpine *X. lorezi*, we observed genetic distinctiveness of both Alps and Fennoscandian populations in all the three phylogenetic approaches. The support for these two populations as separate clusters was high in all three cases. Trees from all the three approaches supported the close phylogenetic relatedness between the Fennoscandian population and two individuals belonging to the NA population with varying degrees of support values, but the phylogenetic position of those individuals was different in the three approaches. We also observed a relatively high F_ST_ value of 0.346 between the Alps and Fennoscandian populations (Table 2).

### Species delimitation

The -log(likelihood) values for all the models tested in tr2 for each of the ten study species can be found in supplementary file. In general, we observed that SODA analysis using a guide tree tended to split the individuals into more species than the one with mapping or the tr2 analyses. The only exception to this pattern was *X. speciosa*, where all three different analyses suggested the same delimitation, i.e., Alps population as one species and Fennoscandian and Altai individuals together as another. For species *E. vittaria*, *E. sudetica* and *X. lorezi*, tr2 and SODA analysis with mapping suggested the same delimitation, i.e., separation of Palearctic arctic-alpine and NA populations into distinct species. In *S. hochenwarthi* and *A. glandon*, SODA analysis with mapping lumped all the populations into single species, suggesting the absence of species-level differences. For Holarctic taxa *A. caja*, *C. sororiata*, *C. tullia* and *M. brunneata*, this analysis lumped Fennoscandian and Alps populations together (except in *M. brunneata*, where all three populations except for NA were lumped together)

We observed that tr2 mostly supported geographically separate populations as separate species. Within the intercontinental category, for the species *C. tullia* and *M. brunneata*, tr2 analysis lumped Eurasian populations into a single species and suggested the separation of the NA population into separate species, agreeing with the SODA analysis with mapping. However, in the case of *M. brunneata*, when we included the model where Yukon individuals were lumped together with Eurasian individuals in tr2, it received the best support as compared to the other models.

## DISCUSSION

Although high-throughput sequencing has revolutionized the field of evolutionary biology by enabling the acquisition of genetic data on massive scales, the delimitation of species continues to be based predominantly on single data types (e.g., morphology) and/or small datasets (e.g., a single genetic marker), despite consensus that such approaches are suboptimal (Will et al. 2005; Schlick-Steiner et al. 2010). This has given rise to a variety of different “cultures” in species delimitation across research communities studying different taxonomic groups. For example, in many groups of insects, the assumption that genital characteristics are not only rich in useful characters, but also vary little within species, as genitalia are believed to form a prezygotic isolation mechanism are popular (Shapiro and Porter 1989; Mikkola 1992). While the idea of integrative taxonomy remains popular among taxonomists, it has also been occasionally challenged, as the pleas for entirely DNA based taxonomy indicate (Tautz et al. 2003). That all taxonomically relevant information is encoded in organisms’ DNA is an important justification of this approach, but we find the high scalability and quantifiability an additional strong argument for it.

The gradual differentiation of allopatric populations inevitably implies that there is a long period of ‘grey zone’ during which it is not possible to delimit the populations taxonomically in an unambiguous way (De Queiroz 2007). Allopatry poses a particular challenge for species delimitation as it is difficult to assess whether the observed difference between two species is the result of an absence of opportunities for interbreeding due to the suppressed of gene flow, the adaptation to different ecological and environmental conditions, or both (Coyne & Orr, 2004; Nosil, 2008). Many if not most species have fragmented distribution patterns (Mutanen et al. 2012) and the range of a large number of species extends even across continents. Thus, the problem of species delimitation under allopatry is widespread, while remaining little addressed in taxonomic literature. For example, the number of Holarctic terrestrial arthropod insect species is probably counted in thousands (Langor and Sheffield 2019) as in Lepidoptera alone they are several hundred (Pohl et al., 2018, also own observation). The taxonomy of many Holarctic species has been unstable because the degree of differentiation observed between the allopatric populations is highly variable, rendering their delimitation often challenging (Kerr et al. 2009; Landry et al. 2013). This makes the taxonomic delimitation of allopatric populations largely opinion-based and hence arbitrary.

In our study we used a recently developed target enrichment kit containing a fixed set of protein-coding loci to understand the patterns of intraspecific variation in allopatric populations of ten arctic-alpine and boreo-montane Lepidoptera species of the Palearctic and Holarctic region. We compared these patterns in nuclear genes to the patterns known from the mtDNA barcodes. In about five out of ten cases we observed that the barcode phylogenies reflected similar phylogenetic patterns as those based on nuclear loci (*S. hochenwarthi, X. speciosa, C. tullia, M. brunneata, X. lorezi*), while in rest we observed mitonuclear discordance. In cases of P-distance comparisons, barcode P-distances in general tended to vary greatly between different comparisons, while the variation in target enrichment P-distances between different comparisons remained lower (Fig. 1). This higher variation in barcode distances, or a mitonuclear discordance in general, can largely be explained by how mitochondrial and nuclear genomes diverge, especially during the fragmentation of larger populations into smaller, geographically isolated ones. First, mitochondrial genomes tend to evolve faster than nuclear genomes due to higher mutation rate as well as selection since the whole mitochondrial genome forms a single linkage group due to lack of recombination (Ballard and Whitlock 2004; Moritz and Cicero 2004). Second, the fragmentation of initial large populations into smaller ones can lead to differential fixation of mtDNA haplotypes in different geographic areas whereas nuclear diversity remains less affected (Després 2019), because random genetic drift is directly linked to effective population size which is four times less for mtDNA as compared to the nuclear genome (Charlesworth 2009). Third, *Wolbachia*- mediated selective sweeps can lead to the local fixation of associated mtDNA haplotypes; this is explained in more detail in the sections below.

From our analyses of nuclear markers, we found the ASTRAL approach to be providing consistently well-resolved topologies which at least in some details tended to conflict with the concatenation-based ML approach. The normalized quartet scores for ASTRAL trees for each of the species are indicative of high gene tree-species tree discordance, especially in cases such as *A. glandon*, *X. speciosa*, and *M. brunneata* (Fig. 4). The causes of this discordance could be either operational or biological. Operational causes include the presence of high GC content and locus informativeness (Espeland et al. 2018). The former can be ruled out as we had removed loci with high GC content (more than 60%) from out dataset. As for the latter one, we can see that in cases such as *E. sudetica*, *X. lorezi*, and *A. caja*, the ML tree showed patterns similar to ASTRAL, which indicates the presence of sufficient phylogenetic signal to recover genetically distinct lineages that belong to the same species. As multispecies coalescent approaches such as ASTRAL takes into account incomplete lineage sorting (ILS) while inferring phylogenies, they are likely to perform better where ILS is significant. Therefore, the discordance between coalescent and concatenation approaches observed in our study can be explained by the likely existence of gene flow due to past contact between the populations especially within the continent, until recent times.

### Within-continental patterns of diversity in Lepidoptera

The patterns of intraspecific variation tended to vary for species in the intra-continental category and no single consistent pattern emerged. In *X. speciosa*, the genetic distinctiveness of the Alps population is weakly supported by phylogenetic approaches as well as F_ST_. Also, the target enrichment P-distances did not show any significant differences between different comparisons. This apparent lack of strong differentiation in the nuclear genome supports the hypothesis that this species had a continuous distribution during the last ice ages (Schmitt and Hewitt 2004). However, the barcode P-distances and structure strongly supports the Alps population as belonging to a separate genetic cluster. Species delimitation analyses likewise suggested that the Alps population is a separate species and lumped Fennoscandian and Altai populations together as one species, suggesting the ongoing divergence of these populations which has not yet reached the completion stage. In *S. hochenwarthi*, the genetic distinctiveness of the Fennoscandian population from that of Alps is strongly supported by the analyses using nuclear genes, but only weakly in the barcodes. In both these species, although similar patterns were observed in phylogenies for barcodes and nuclear markers, P-distance comparisons using barcodes varied greatly demonstrating that DNA barcodes may not always reliably reflect the true intraspecific variation.

We observed an arctic-alpine separation based on analyses of nuclear markers in 3 out of 10 analyzed species – *E. vittaria, E. sudetica,* and *X. lorezi* – that were supported by our species delimitation analyses. In the seven other cases, we did not observe clear genetic separation between the Alps and Fennoscandian populations. In *E. vittaria*, we observed rather inconsistent, weakly supported relationships from our phylogenetic analyses. The most straightforward reason for this could be lack of sufficient phylogenetic signal from the selected nuclear genes to delimit the intraspecific variation. It is also likely that the gene flow has ceased since the last Glacial Period and the gradual divergence of the populations from two regions is underway but has not yet reached completion, hence providing an excellent example of the “grey zone” and inherently arbitrary nature of delineation of such populations into species. In the P-distance comparison, barcode distances supported deep divergence between Alps and Fennoscandia (median value of P-distances more than 5%), and previous studies have revealed even higher divergence (6.47%, Mutanen et al. 2012). These deep intraspecific barcode divergences can arise as a result of *Wolbachia*-mediated mitochondrial introgression and subsequent selective sweeps of the mitochondrial genome (Hilgenboecker et al. 2008), although we do not know the potential “donator” species and which of the two populations has undergone such a selective sweep. However, the separation of the populations is also reflected in the nuclear markers, where the median target enrichment P-distance between Fennoscandia and Alps was slightly higher than 1%, suggesting that further assessment of the taxonomic status is required.

In *E. sudetica*, monophyly of Fennoscandian population was supported from all three phylogenies. The Pyrenees individuals show a striking divergence especially from PCA and P-distances and in the ASTRAL tree. STRUCTURE, on the other hand, did not support the genetic separation between Alps and Pyrenees populations, but supported the genetic distinctness of the Apennines individual. Species delimitation analyses further supported the three populations as separate species, thus at least one of the three populations analyzed likely belongs to an unrecognized cryptic species, requiring further taxonomic revisional work. For *X. lorezi*, a significant separation of arctic-alpine populations was supported by phylogenies as well as barcode P-distances, F_ST_ and species delimitation. We could not analyze the Nearctic population due to insufficient sampling, but our results point to it being related to the Fennoscandian population.

It is likely that the populations from Fennoscandia and the European mountains such as Alps, Pyrenees and Apennines are descendants of large populations that have dwelled in the periglacial tundra belt during the last glaciations (Schmitt et al. 2010). Most often, separations in arctic-alpine species between the Northern populations and the high mountain systems in central and Southern Europe (often the Alps) are recent (i.e., post-glacial) phenomena, which is the result of species retreating northwards and into the mountain systems in the south during post-glacial periods (Schmitt et al. 2010). On the other hand, species such as *X. speciosa* which do not show significant separation of populations, were likely widely distributed in the cold steppe belt during glaciation, enabling a permanent gene-flow over the continent/within continent (Varga & Schmitt, 2008).

### Patterns of diversity across the continents in Lepidoptera

We studied genetic patterns of five Holarctic species in light of target enrichment data set and compared these patterns to those observed in their DNA barcodes. Holarctic distributions are often the result of either past land connections over the Pacific (Beringian) or of dispersal (Mikkola et al. 1991), and species presently occurring in both continents may therefore have different biogeographical histories. In *A. glandon*, a rather weak differentiation of the NA population in the nuclear data and no differentiation in the barcodes was observed. Species delimitation analyses using tr2 inferred the Palearctic and Nearctic populations as separate species while SODA analysis with mapping suggested the presence of a single species. This mitonuclear discordance, as well as the lack of phylogenetic resolution in the phylogenomic ML tree, points to incomplete lineage sorting (as discussed above) and absence of species level differences in the populations of *A. glandon*.

In *A. caja*, the Nearctic population was revealed to be differentiated from the Palearctic counterpart in the target enrichment data, but DNA barcodes did not reveal any differentiation between the continents. The Nearctic population was further observed to be separated into two clades. This separation is also evident from the P-distance boxplots, where the Nearctic population displays high variation in within-population divergences. We also observed high F_ST_ values in *A. caja* for NA-FEN and NA-Alps comparisons, while slightly lower values for the NA-Altai comparison. This coincides with our STRUCTURE observations where signs of gene exchange were detected between Nearctic and Altai populations, which could be the result of past contact between two populations through the Beringian land bridge. Tr2 analysis split the geographically separate populations into distinct species, which is possibly due to the presence of isolation by distance effect, as IBD was revealed to be significant in the case of *A. caja*. Similar reasoning might hold true for *C. sororiata*, where tr2 suggested geographically distinct populations as separate species, where otherwise no genetic differentiation was observed. In the case of *C. tullia*, the presence of the Eurasian gene pool in Nunavut suggests the possible dispersal of this species across the Beringian/Pacific from far eastern Eurasia. A detailed study including more samples from this region would be needed.

*Carsia sororiata* and *M. brunneata* are examples of Holarctic species with a boreal distribution whose larvae feed on Holarctic *Vaccinium*. In the case of *M. brunneata*, the presence of Eurasian gene pool in two individuals from Yukon suggests historical contact between this region and Eurasia, as Alaska and Yukon were effectively a part of the Palearctic region connected to northeastern Siberia through the Beringian land bridge, which was present until *c.* 15,000 years ago (Mikkola et al. 1991; Brubaker et al. 2005). For *M. brunneata*, the Nearctic population (except for Yukon individuals) is suggested as a different species by tr2 analyses. It is difficult to assess whether this is due to isolation by distance effect or if it reflects the real divergence of samples from the Eastern Nearctic. The apparent lack of differentiation of any population in *C. sororiata* suggests that it had a widespread distribution even during glaciation, thus the species-level differences might be absent from geographically separated populations. However, although phylogenies failed to reveal differentiation in the case of *C. sororiata* populations, the high F_ST_ values between NA-Alps and Alps-Altai indicates the presence of genetic differentiation in the Alps population, possibly after the glaciation period.

We observed that our SODA results where we used ASTRAL tree as a guide mostly coincided with the ASTRAL tree structure (Figure 4a-j), possibly due to SODA taking gene tree topologies into account while computing delimitation. While delimitation programs such as SODA are considerably faster than the more commonly used methods such as BPP, they are also prone to errors. As the accuracy of this method depends on input gene trees, errors in gene tree tree-topologies may bias SODA towards over-splitting (Patel et al. 2013; Mirarab et al. 2014). SODA is also known to be sensitive to population structure within species (Rabiee and Mirarab 2020). These two factors could explain the observed over-splitting of populations into species in several cases.

### Towards consistent delimitation of allopatric populations – Can standard nuclear loci solve the problem?

A previous study has suggested a genetic divergence threshold between 0.5% to 2% in the nuclear genomic data as typical for taxa belonging to a grey zone, i.e. falling in the interface between intra- and interspecific variation (Roux et al. 2016). Another study found that a 0% to 1.1% divergence in genome-wide RADseq data represented within-species polymorphism/intraspecific variability (Cariou et al. 2020). In our case, *C. tullia* is the only Holarctic species that showed P-distance divergence of higher than 1% between the continents in target enrichment data, suggesting genetic differentiation at species level and not just the isolation by distance effect. However, there is no well-established and objective threshold to distinguish intraspecific variation from interspecific variation in the nuclear genome and the thresholds of pairwise genetic distances for mitochondrial DNA have shown to be inconsistent and overlapping between intra- and interspecific comparisons (Meier et al. 2006). In *A. caja* and *M. brunneata*, we found the NA population to be genetically distinct in trees inferred using several hundred nuclear loci, but the P-distance between continents based on the same data was low (<1%). This could point to subspecies level differentiation which is mainly the effect of geographic isolation.

Hey & Pinho, (2012) identified a threshold value of F_ST_ = 0.35 – to coincide with the current taxonomic status the most. Above this value the entities were identified as species, and below as populations. Thus, based on F_ST_ alone, the Nearctic population of *A. caja* deserves distinct species status, as do the Alps and Fennoscandian populations in *X. lorezi* and *E. sudetica*. However, several studies have found a significant statistical association of F_ST_ values with divergence times (Lessios and Robertson 2006; Marko and Hart 2012), thus the low F_ST_ values might not necessarily point to low genetic differentiation, but rather might be the result of the recent divergences (and vice a versa for high F_ST_ values).

Given the inherent arbitrariness, will it be possible to create universal standards for the delimitation of allopatric populations? Would this be a desirable goal at all? Our response is “Yes”, although we acknowledge that it will likely be very difficult to agree about those standards among the taxonomic community. We find that among all possible solutions, the prevailing situation, i.e., full lack of an epistemological solution to the allopatry conundrum, is the least optimal situation, at least in terms of consistency and stability. As the delimitation of allopatric populations is bound to be based on arbitrary criteria, applying the same principles throughout taxa, or at least very broadly, would be an optimal solution. We further find that using a large set of universal genetic markers, such as the USCO markers as proposed by Eberle et al. (2020), and implemented by Dietz et al. (2022) would provide the best means for that. The benefits of this approach would be multiple, but at its best we hope that it would reduce the subjectivity of types of traits used and hence critically improve the scalability and quantifiability of the taxonomic judgement. Our study demonstrates that mitochondrial markers may provide only an indication of the genetic distinctiveness of populations and that mitonuclear discordance is a widespread phenomenon. The same is likely true for any single genetic marker as the genealogy of any single gene may deviate from that of a species (Pamilo and Nei 1988; Maddison 1997).

## CONCLUSIONS

Our study is among the first to focus on species delimitation on a very broad geographic scale and based on genome-wide data. After witnessing a large number of taxa with various and often fully unexplained patterns of divergences observed in DNA barcode data sets across almost all animal groups (e.g., Huemer et al., 2018; Mutanen et al., 2012; Roe & Sperling, 2007; Shearer & Coffroth, 2008; Ward, 2009), we sought to gain a genomic insight on these problems. These patterns are presently poorly understood and desperately call for explanations. We find that such in-depth insights into these patterns are best obtained by genomics approaches. Our study demonstrates that patterns observed in mitochondrial DNA may have a variety of biological explanations and may not always reflect true genealogy, suggesting that multiple genetic markers are required to draw an accurate picture of genetic differences between populations. Modern high-throughput DNA sequencing tools, like those employed here, have significant potential to provide a solid foundation for species delimitation, and might critically reduce the subjectivity and non-scalability of the delimitation process. We therefore propose that as delimitation of allopatric populations cannot be made in any objective way, it would be best if based on quantifiable data and standard criteria, even if arbitrary. The recently proposed DNA taxonomy scheme (Eberle et al. 2020) applied to the delimitation of allopatric populations would provide a solid foundation in terms of quantifiability. Finally, we acknowledge that our analysis suffers from limited sampling and hence may not provide an accurate picture of taxonomic status of the studied taxa. Consequently, we refrain to make any formal taxonomic actions here. For example, our underlying assumption of the uniformity of the North American populations as single entities is often false as these populations frequently show strong intra-continental sub-structuring (D’Ercole et al. 2021; Campbell et al. 2022).

## DATA ACCESSIBILITY

All raw cleaned sequence data are archived on the NCBI Sequence Read Archive under BioProject ID PRJNA922447. Alignments and other unpublished data are available on the dryad via link https://datadryad.org/stash/share/PPtY7vmqqL4-IHte0Arx43Ct_rvoZ687zhnbtG0VzM4 (doi: 10.5061/dryad.ht76hdrm0)

## FUNDING

MM was funded by the Academy of Finland (grant no. 314702). JRd was supported through the University of Guelph’s ‘Food From Thought’ project, funded by the Canada First Research Excellence Fund. MJ would like to thank University of Oulu scholarship Foundation and Oulun Luonnonystäväin Yhdistys ry for providing funding to continue her doctoral studies.

## ACKNOWLEDGEMENTS

We would like to thank Laura Törmälä for her valuable help with the target enrichment laboratory work and Hannele Parkkinen for help with the barcoding. We would also like to thank CSC Finland for providing computational resources to carry out bioinformatics analyses, and CSC staff for their help with bioinformatics troubleshooting.

## Notes

### Competing Interest Statement

The authors have declared no competing interest.

## REFERENCES

Ballard J.W.O., Whitlock M.C. 2004. The incomplete natural history of mitochondria. Mol. Ecol. 13:729–744.

Bergsten J., Bilton D.T., Fujisawa T., Elliott M., Monaghan M.T., Balke M., Hendrich L., Geijer J., Herrmann J., Foster G.N., Ribera I., Nilsson A.N., Barraclough T.G., Vogler A.P. 2012. The effect of geographical scale of sampling on DNA barcoding. Syst. Biol. 61:851–869.

Breinholt J.W., Earl C., Lemmon A.R., Lemmon E.M., Xiao L., Kawahara A.Y. 2018. Resolving relationships among the megadiverse butterflies and moths with a novel pipeline for anchored phylogenomics. Syst. Biol. 67:78–93.

Brubaker L.B., Anderson P.M., Edwards M.E., Lozhkin A.V. 2005. Beringia as a glacial refugium for boreal trees and shrubs: new perspectives from mapped pollen data. J. Biogeogr. 32:833–848.

Campbell E.O., MacDonald Z.G., Gage E.V., Gage R.V., Sperling F.A.H. 2022. Genomics and ecological modelling clarify species integrity in a confusing group of butterflies. Mol. Ecol. 31:2400–2417.

Cariou M., Henri H., Martinez S., Duret L., Charlat S. 2020. How consistent is RAD-seq divergence with DNA-barcode based clustering in insects? Mol. Ecol. Resour. 20:1294–1298.

Charlesworth B. 2009. Effective population size and patterns of molecular evolution and variation. Nat Rev Genet. 10:195–205.

Chen S., Zhou Y., Chen Y., Gu J. 2018. fastp: an ultra-fast all-in-one FASTQ preprocessor. Bioinformatics. 34:i884–i890.

Chernomor O., Von Haeseler A., Minh B.Q. 2016. Terrace aware data structure for phylogenomic inference from supermatrices. Syst. Biol. 65:997–1008.

Coyne J.A., Orr H.A. 2004. Speciation. Oxford, New York: Oxford University Press.

Cummings M.P. 2004. PAUP* [Phylogenetic Analysis Using Parsimony (and Other Methods)]. Dictionary of Bioinformatics and Computational Biology. John Wiley & Sons, Ltd.

Dayrat B. 2005. Towards integrative taxonomy. Biol. J. Linn. Soc. 85:407–417.

De Queiroz K. 2007. Species concepts and species delimitation. Syst. Biol. 56:879–886.

D’Ercole J., Dincă V., Opler P.A., Kondla N., Schmidt C., Phillips J.D., Robbins R., Burns J.M., Miller S.E., Grishin N., Zakharov E.V., DeWaard J.R., Ratnasingham S., Hebert P.D.N. 2021. A DNA barcode library for the butterflies of North America. PeerJ. 9:e11157.

Després L. 2019. One, two or more species? Mitonuclear discordance and species delimitation. Mol. Ecol. 28:3845–3847.

Dietz L., Eberle J., Mayer C., Kukowka S., Bohacz C., Baur H., Espeland M., Huber B.A., Hutter C., Mengual X., Peters R.S., Vences M., Wesener T., Willmott K., Misof B., Niehuis O., Ahrens D. 2022. Standardized nuclear markers improve and homogenize species delimitation in Metazoa. Methods Ecol. Evol. 00:1–13.

Earl D.A., vonHoldt B.M. 2012. STRUCTURE HARVESTER: A website and program for visualizing STRUCTURE output and implementing the Evanno method. Conserv. Genet. Resour. 4:359–361.

Eberle J., Ahrens D., Mayer C., Niehuis O., Misof B. 2020. A plea for standardized nuclear markers in metazoan DNA taxonomy. Trends Ecol. Evol. 35:336–345.

Edgar R.C. 2004. MUSCLE: a multiple sequence alignment method with reduced time and space complexity. BMC Bioinf. 5:113.

Erickson K.L., Pentico A., Quattrini A.M., McFadden C.S. 2020. New approaches to species delimitation and population structure of anthozoans: Two case studies of octocorals using ultraconserved elements and exons. Mol. Ecol. Resour. 00:1–15.

Espeland M., Breinholt J., Willmott K.R., Warren A.D., Vila R., Toussaint E.F.A., Maunsell S.C., Aduse-Poku K., Talavera G., Eastwood R., Jarzyna M.A., Guralnick R., Lohman D.J., Pierce N.E., Kawahara A.Y. 2018. A comprehensive and dated phylogenomic analysis of butterflies. Curr. Biol. 28:770–778.e5.

Evanno G., Regnaut S., Goudet J. 2005. Detecting the number of clusters of individuals using the software STRUCTURE: A simulation study. Mol. Ecol. 14:2611–2620.

Fujisawa T., Aswad A., Barraclough T.G. 2016. A rapid and scalable method for multilocus species delimitation using bayesian model comparison and rooted triplets. Syst. Biol. 65:759–771.

Funk D.J., Omland K.E. 2003. Species-Level paraphyly and polyphyly: Frequency, causes, and consequences, with insights from animal mitochondrial DNA. Annu. Rev. Ecol. Evol. Syst. 34:397–423.

Godfray H.C.J. 2002. Challenges for taxonomy. Nature. 417:17–19.

Goudet J. 2005. HIERFSTAT, a package for R to compute and test hierarchical F-statistics. Mol. Ecol. Notes. 5:184–186.

Grabherr M.G., Haas B.J., Yassour M., Levin J.Z., Thompson D.A., Amit I., Adiconis X., Fan L., Raychowdhury R., Zeng Q., Chen Z., Mauceli E., Hacohen N., Gnirke A., Rhind N., di Palma F., Birren B.W., Nusbaum C., Lindblad-Toh K., Friedman N., Regev A. 2011. Full-length transcriptome assembly from RNA-Seq data without a reference genome. Nat. Biotechnol. 29:644–652.

Grummer J.A., Morando M.M., Avila L.J., Sites J.W., Leaché A.D. 2018. Phylogenomic evidence for a recent and rapid radiation of lizards in the Patagonian *Liolaemus fitzingerii* species group. Mol. Phylogenet. Evol. 125:243–254.

Guindon S., Dufayard J.F., Lefort V., Anisimova M., Hordijk W., Gascuel O. 2010. New algorithms and methods to estimate maximum-likelihood phylogenies: Assessing the performance of PhyML 3.0. Syst. Biol. 59:307–321.

György Z., Tóth E.G., Incze N., Molnár B., Höhn M. 2018. Intercontinental migration pattern and genetic differentiation of arctic-alpine *Rhodiola rosea* L.: A chloroplast DNA survey. Ecol. Evol. 8:11508–11521.

Herrera S., Shank T.M. 2016. RAD sequencing enables unprecedented phylogenetic resolution and objective species delimitation in recalcitrant divergent taxa. Mol. Phylogenet. Evol. 100:70–79.

Hewitt G.M. 2004. Genetic consequences of climatic oscillations in the Quaternary. Philos. Trans. R. Soc. London, Ser. B. 359:183–195.

Hey J., Pinho C. 2012. Population genetics and objectivity in species diagnosis. Evolution. 66:1413–1429.

Hilgenboecker K., Hammerstein P., Schlattmann P., Telschow A., Werren J.H. 2008. How many species are infected with Wolbachia? – a statistical analysis of current data. FEMS Microbiol. Lett. 281:215–220.

Hoang D.T., Chernomor O., Von Haeseler A., Minh B.Q., Vinh L.S. 2018. UFBoot2: Improving the ultrafast bootstrap approximation. Mol. Biol. Evol. 35:518–522.

Hou Y., Nowak M.D., Mirré V., Bjorå C.S., Brochmann C., Popp M. 2015. Thousands of RAD-seq loci fully resolve the phylogeny of the highly disjunct arctic-alpine genus *Diapensia* (Diapensiaceae). PLoS ONE. 10.

Huemer P., Hebert P.D.N., Mutanen M., Wieser C., Wiesmair B., Hausmann A., Yakovlev R., Möst M., Gottsberger B., Strutzenberger P., Fiedler K. 2018. Large geographic distance versus small DNA barcode divergence: Insights from a comparison of European to South Siberian Lepidoptera. PloS ONE. 13:e0206668.

Huemer P., Mutanen M., Sefc K.M., Hebert P.D.N. 2014. Testing DNA barcode performance in 1000 species of European Lepidoptera: Large geographic distances have small genetic impacts. PLoS ONE. 9:1–21.

Jackson N.D., Carstens B.C., Morales A.E., O’Meara B.C. 2017. Species delimitation with gene flow. Syst. Biol. 66:799–812.

Jombart T., Ahmed I. 2011. adegenet 1.3-1: new tools for the analysis of genome-wide SNP data. Bioinformatics. 27:3070–3071.

Junier T., Zdobnov E.M. 2010. The Newick utilities: high-throughput phylogenetic tree processing in the Unix shell. Bioinformatics. 26:1669–1670.

Kalyaanamoorthy S., Minh B.Q., Wong T.K.F., Von Haeseler A., Jermiin L.S. 2017. ModelFinder: fast model selection for accurate phylogenetic estimates. Nat. Methods. 14:587–589.

Kawahara A. Y., & Breinholt J. W. 2014. Phylogenomics provides strong evidence for relationships of butterflies and moths. Proc. R. Soc. B. 281:1788. https://doi.org/10.1098/rspb.2014.0970

Kerr K.C., Birks S.M., Kalyakin M.V., Red’kin Y.A., Koblik E.A., Hebert P.D. 2009. Filling the gap - COI barcode resolution in eastern Palearctic birds. Front. Zool. 6:29.

Knowlton N. 1993. Sibling species in the sea. Annu. Rev. Ecol. Syst. 24:189–216.

Kopelman N.M., Mayzel J., Jakobsson M., Rosenberg N.A., Mayrose I. 2015. Clumpak: a program for identifying clustering modes and packaging population structure inferences across K. Mol. Ecol. Resour. 15:1179–1191.

Landry J.-F., Nazari V., Dewaard J.R., Mutanen M., Lopez-Vaamonde C., Huemer P., Hebert P.D.N. 2013. Shared but overlooked: 30 species of Holarctic Microlepidoptera revealed by DNA barcodes and morphology. Zootaxa. 3749:1–93.

Lanfear R., Calcott B., Kainer D., Mayer C., Stamatakis A. 2014. Selecting optimal partitioning schemes for phylogenomic datasets. BMC Evol. Biol. 14:1–14.

Langor D.W., Sheffield C.S. 2019. The Biota of Canada: Terrestrial Arthropods. ZooKeys. 819:1–4.

Leaché A.D., Fujita M.K., Minin V.N., Bouckaert R.R. 2014. Species delimitation using genome-wide SNP Data. Syst. Biol. 63:534–542.

Lessios H.A., Robertson D.R. 2006. Crossing the impassable: genetic connections in 20 reef fishes across the eastern Pacific barrier. Proc. Biol. Sci. 273:2201–2208.

Lindholm M., d’Auriac M.A., Thaulow J., Hobæk A. 2016. Dancing around the pole: holarctic phylogeography of the Arctic fairy shrimp *Branchinecta paludosa* (Anostraca, Branchiopoda). Hydrobiologia. 772:189–205.

Lischer H.E.L., Excoffier L. 2012. PGDSpider: An automated data conversion tool for connecting population genetics and genomics programs. Bioinformatics. 28:298–299.

Maddison W.P. 1997. Gene trees in species trees. Syst. Biol. 46:523–536.

Mai U., Mirarab S. 2018. TreeShrink: fast and accurate detection of outlier long branches in collections of phylogenetic trees. BMC Genomics. 19:272.

Mantel N. 1967. The detection of disease clustering and a generalized regression approach. Cancer Res. 27:209–220.

Marko P.B., Hart M.W. 2012. Retrospective coalescent methods and the reconstruction of metapopulation histories in the sea. Evol. Ecol. 26:291–315.

Mayer C., Dietz L., Call E., Kukowka S., Martin S., Espeland M. 2021. Adding leaves to the Lepidoptera tree: capturing hundreds of nuclear genes from old museum specimens. Syst. Entomol. 46:649–671.

Meier R., Shiyang K., Vaidya G., Ng P.K.L. 2006. DNA barcoding and taxonomy in Diptera: A tale of high intraspecific variability and low identification success. Syst. Biol. 55:715–728.

Meyer C.P., Paulay G. 2005. DNA barcoding: Error rates based on comprehensive sampling. PLoS Biol. 3:1–10.

Mikkola K. 1992. Evidence for lock-and-key mechanisms in the internal genitalia of the *Apamea* moths (Lepidoptera, Noctuidae). Syst. Entomol. 17:145–153.

Mikkola K., Lafontaine D., Kononenko V. 1991. Zoogeography of the Holarctic species of the Noctuidae (Lepidoptera): importance of the Beringian refuge. Entomologica Fennica. 2:157–173.

Minh B.Q., Schmidt H.A., Chernomor O., Schrempf D., Woodhams M.D., Von Haeseler A., Lanfear R., Teeling E. 2020. IQ-TREE 2: New models and efficient methods for phylogenetic inference in the genomic era. Mol. Biol. Evol. 37:1530–1534.

Mirarab S., Bayzid Md.S., Boussau B., Warnow T. 2014. Statistical binning enables an accurate coalescent-based estimation of the avian tree. Science. 346:1250463.

Misof B., Misof K. 2009. A Monte Carlo approach successfully identifies randomness in multiple sequence alignments: A more objective means of data exclusion. Syst. Biol. 58:21–34.

Moritz C., Cicero C. 2004. DNA barcoding: Promise and pitfalls. PLoS Biol. 2.

Muster C., Berendonk T.U. 2006. Divergence and diversity: Lessons from an arctic-alpine distribution (*Pardosa saltuaria* group, Lycosidae). Mol. Ecol. 15:2921–2933.

Mutanen M., Hausmann A., Hebert P.D.N., Landry J.F., de Waard J.R., Huemer P. 2012. Allopatry as a gordian knot for taxonomists: Patterns of DNA barcode divergence in arctic-alpine Lepidoptera. PLoS ONE. 7.

Mutanen M., Kivelä S.M., Vos R.A., Doorenweerd C., Ratnasingham S., Hausmann A., Huemer P., Dincă V., van Nieukerken E.J., Lopez-Vaamonde C., Vila R., Aarvik L., Decaëns T., Efetov K.A., Hebert P.D.N., Johnsen A., Karsholt O., Pentinsaari M., Rougerie R., Segerer A., Tarmann G., Zahiri R., Godfray H.C.J. 2016. Species-level para- and polyphyly in DNA barcode gene trees: Strong operational bias in European Lepidoptera. Syst. Biol. 65:1024–1040.

Nosil P. 2008. Ernst Mayr and the integration of geographic and ecological factors in speciation. Biol. J. Linn. Soc. 95:26–46.

Packer L., Gibbs J., Sheffield C., Hanner R. 2009. DNA barcoding and the mediocrity of morphology. Mol. Ecol. Resour. 9:42–50.

Padial J.M., Miralles A., De La Riva I., Vences M. 2010. The integrative future of taxonomy. Front. Zool. 7:16.

Pamilo P., Nei M. 1988. Relationships between gene trees and species trees. Mol. Biol. Evol. 5:568–583.

Pante E., Schoelinck C., Puillandre N. 2015. From integrative taxonomy to species description: One step beyond. Syst. Biol. 64:152–160.

Patel S., Kimball R.T., Braun E.L. 2013. Error in phylogenetic estimation for bushes in the tree of life. J. Phylogen. Evolution. Biol. 1:1–10.

Petersen M., Meusemann K., Donath A., Dowling D., Liu S., Peters R.S., Podsiadlowski L., Vasilikopoulos A., Zhou X., Misof B., Niehuis O. 2017. Orthograph: a versatile tool for mapping coding nucleotide sequences to clusters of orthologous genes. BMC Bioinf. 18:111.

Pie M.R., Bornschein M.R., Ribeiro L.F., Faircloth B.C., McCormack J.E. 2019. Phylogenomic species delimitation in microendemic frogs of the Brazilian Atlantic Forest. Mol. Phylogenet. Evol. 141:106627.

Pohl G.R., Landry J.-F., Schmidt B.C., Lafontaine J.D., Troubridge J.T., Macaulay A.D., Van Nieukerken E.J., deWaard J.R., Dombroskie J.J., Klymko J., Nazari V., Stead K. 2018. Annotated checklist of the moths and butterflies (Lepidoptera) of Canada and Alaska.

Pritchard J.K., Stephens M., Donnelly P. 2000. Inference of population structure using multilocus genotype data. Genetics. 155:945–959.

Rabiee M., Mirarab S. 2020. SODA: multi-locus species delimitation using quartet frequencies. Bioinformatics. 36:5623–5631.

Roe A.D., Sperling F.A.H. 2007. Patterns of evolution of mitochondrial cytochrome c oxidase I and II DNA and implications for DNA barcoding. Mol. Phylogenet. Evol. 44:325–345.

Roux C., Fraïsse C., Romiguier J., Anciaux Y., Galtier N., Bierne N. 2016. Shedding light on the grey zone of speciation along a continuum of genomic divergence. PLOS Biol. 14:e2000234.

Schlick-Steiner B.C., Steiner F.M., Seifert B., Stauffer C., Christian E., Crozier R.H. 2010. Integrative taxonomy: A multisource approach to exploring biodiversity. Annu. Rev. of Entomol. 55:421–438.

Schmitt T. 2007. Molecular biogeography of Europe: Pleistocene cycles and postglacial trends. Front. Zool. 4.

Schmitt T., Hewitt G.M. 2004. Molecular biogeography of the arctic-alpine disjunct burnet moth species *Zygaena exulans* (Zygaenidae, Lepidoptera) in the Pyrenees and Alps. J. Biogeogr. 31:885–893.

Schmitt T., Hewitt G.M., Müller P. 2006. Disjunct distributions during glacial and interglacial periods in mountain butterflies: *Erebia epiphron* as an example. J. Evol. Biol. 19:108–113.

Schmitt T., Louy D., Zimmermann E., Habel J.C. 2016. Species radiation in the Alps: multiple range shifts caused diversification in Ringlet butterflies in the European high mountains. Org. Divers. Evol. 16:791–808.

Schmitt T., Muster C., Schönswetter P. 2010. Are disjunct alpine and arctic-alpine animal and plant species in the Western Palearctic really relics of a cold past? Relict Species: Phylogeography and Conservation Biology. Springer-Verlag Berlin Heidelberg. p. 239–252.

Shapiro A.M., Porter A.H. 1989. The lock-and-key hypothesis: Evolutionary and biosystematic interpretation of insect genitalia. Annu. Rev. Entomol. 34:231–245.

Shearer T.L., Coffroth M.A. 2008. DNA BARCODING: Barcoding corals: limited by interspecific divergence, not intraspecific variation. Mol. Ecol. Resour. 8:247–255.

Slater G., Birney E. 2005. Automated generation of heuristics for biological sequence comparison. BMC Bioinf. 6:31.

Tautz D., Arctander P., Minelli A., Thomas R.H., Vogler A.P. 2003. A plea for DNA taxonomy. Trends Ecol. Evol. 18:70–74.

Varga Z.S., Schmitt T. 2008. Types of oreal and oreotundral disjunctions in the western Palearctic. Biol. J. Linn. Soc. 93:415–430.

Ward R.D. 2009. DNA barcode divergence among species and genera of birds and fishes. Mol. Ecol. Resour. 9:1077–1085.

Wickham H (2016). ggplot2: Elegant Graphics for Data Analysis. Springer-Verlag New York. ISBN 978-3-319-24277-4, http://ggplot2.org.

Wiemers M., Fiedler K. 2007. Does the DNA barcoding gap exist? – a case study in blue butterflies (Lepidoptera: Lycaenidae). Front. Zool. 4:8.

Will K.W., Mishler B.D., Wheeler Q.D. 2005. The perils of DNA barcoding and the need for integrative taxonomy. Syst. Biol. 54:844–851.

Yang Z. 2015. The BPP program for species tree estimation and species delimitation. Curr. Zool. 61:854–865.

Yeates D.K., Seago A., Nelson L., Cameron S.L., Joseph L., Trueman J.W.H. 2011. Integrative taxonomy, or iterative taxonomy? Syst. Entomol. 36:209–217.

Zarza E., Faircloth B.C., Tsai W.L.E., Bryson R.W., Klicka J., McCormack J.E. 2016. Hidden histories of gene flow in highland birds revealed with genomic markers. Mol. Ecol. 25:5144–5157.

Zhang C., Rabiee M., Sayyari E., Mirarab S. 2018. ASTRAL-III: Polynomial time species tree reconstruction from partially resolved gene trees. BMC Bioinf. 19:15–30.

