## Supplementary material for "Species Delimitation Under Allopatry: Genomic Divergences Within and Across Continents in Lepidoptera"

**Table S1:** Metadata for specimens analyzed

| Sample ID | SRA  accession | Species | Country | Locality | Lat | Lon | No. informative loci | % missing data |
| --- | --- | --- | --- | --- | --- | --- | --- | --- |
| TLMF Lep 02482 | SAMN32653148 | *E. vittaria* | Italy | Ritten/ Obergruenwald | 46.59 N | 11.44 E | 434 | 18,56 % |
| TLMF Lep 02417 | SAMN32653149 | *E. vittaria* | Italy | Ritten/ Obergruenwald | 46.59 N | 11.44 E | 417 | 21,80 % |
| TLMF Lep 16050 | SAMN32653150 | *E. vittaria* | Austria | Ellbachtal, unterer Kaiserboden | 47.54 N | 11.93 E | 423 | 18,85 % |
| TLMF Lep 00248 | SAMN32653151 | *E. vittaria* | Austria | Petzen N, Obere Krischa | 46.51 N | 14.76 E | 444 | 15,31 % |
| TLMF Lep 30328 | SAMN32653152 | *E. vittaria* | Liechtenstein | S Falleck | 49.09 N | 9.35 E | 431 | 16,28 % |
| TLMF Lep 30329 | SAMN32653153 | *E. vittaria* | Liechtenstein | S Falleck | 49.09 N | 9.35 E | 431 | 16,23 % |
| TLMF Lep 30330 | SAMN32653154 | *E. vittaria* | Austria | Fliess/ Pillermoor | 47.07 N | 10.39 E | 455 | 12,06 % |
| TLMF Lep 30331 | SAMN32653155 | *E. vittaria* | Austria | Fliess/ Pillermoor | 47.07 N | 10.39 E | 456 | 11,55 % |
| MM06327 | SAMN32653156 | *E. vittaria* | Finland |  |  |  | 321 | 36,67 % |
| MM27376 | SAMN32653157 | *E. vittaria* | Finland |  | 67.79 N | 29.72 E | 427 | 17,62 % |
| MM27377 | SAMN32653158 | *E. vittaria* | Finland | Kiiminki, Ostrobottnia ouluensis |  |  | 480 | 7,95 % |
| MM27378 | SAMN32653159 | *E. vittaria* | Finland | Kiiminki, Ostrobottnia ouluensis |  |  | 474 | 9,74 % |
| TLMF Lep 00128 | SAMN32653160 | *E. sudetica* | Italy |  | 44.39 N | 7.12 E | 449 | 24,63 % |
| TLMF Lep 22067 | SAMN32653161 | *E. sudetica* | Italy | Umg. Franzenshoehe | 46.53 N | 10.48 E | 495 | 18,25 % |
| TLMF Lep 22066 | SAMN32653162 | *E. sudetica* | Italy | Umg. Franzenshoehe | 46.53 N | 10.48 E | 427 | 29,99 % |
| TLMF Lep 22068 | SAMN32653163 | *E. sudetica* | Andorra | Port de Cabus | 42.55 N | 1.42 E | 453 | 25,29 % |
| TLMF Lep 22069 | SAMN32653164 | *E. sudetica* | Andorra | Port de Cabus | 42.55 N | 1.42 E | 455 | 22,70 % |
| TLMF Lep 03039 | SAMN32653165 | *E. sudetica* | Italy | Monte Terminillo N | 42.48 N | 13.01 E | 436 | 26,74 % |
| TLMF Lep 30321 | SAMN32653166 | *E. sudetica* | Italy | Prov. Cuneo Alpi Cozie Demonte NW Colle Valcavera NE | 44.23 N | 7.62 E | 448 | 22,69 % |
| TLMF Lep 30322 | SAMN32653167 | *E. sudetica* | Andorra | Port de Cabus | 42.32 N | 1.25 E | 462 | 20,93 % |
| MM27175 | SAMN32653168 | *E. sudetica* | Finland |  |  |  | 417 | 27,89 % |
| MM27176 | SAMN32653169 | *E. sudetica* | Finland |  |  |  | 446 | 24,83 % |
| MM27177 | SAMN32653170 | *E. sudetica* | Finland |  |  |  | 442 | 23,91 % |
| MM25967 | SAMN32653171 | *E. sudetica* | Finland |  |  |  | 485 | 18,99 % |
| MM25969 | SAMN32653172 | *E. sudetica* | Finland |  |  |  | 494 | 18,21 % |
| MM27418 | SAMN32653173 | *E. sudetica* | Finland |  | 62.59 N | 29.56 E | 457 | 22,35 % |
| MM27419 | SAMN32653174 | *E. sudetica* | Finland |  | 59.83 N | 23.01 E | 430 | 27,29 % |
| MM27420 | SAMN32653175 | *E. sudetica* | Finland |  | 59.83 N | 23.01 E | 414 | 27,84 % |
| TLMF Lep 11234 | SAMN32653176 | *S. hochenwarthi* | Italy | Obere Tartscher Alm | 46.54 N | 10.49 E | 556 | 18,04 % |
| TLMF Lep 20671 | SAMN32653177 | *S. hochenwarthi* | Russia | 17 km NNE Kokorya vill., Chikhacheva Mts. Range, Talduair Mt., valley of Sajlyugem river | 50.02 N | 89.23 E | 489 | 27,84 % |
| TLMF Lep 20670 | SAMN32653178 | *S. hochenwarthi* | Russia | Northern part of Ukok plateau, Zhumaly riber basin | 49.50 N | 88.08 E | 479 | 29,54 % |
| TLMF Lep 30332 | SAMN32653179 | *S. hochenwarthi* | Switzerland | Graubuenden Unterengadin Scuol Champatsch | 46.83 N | 10.26 E | 583 | 13,80 % |
| TLMF Lep 30310 | SAMN32653180 | *S. hochenwarthi* | Switzerland | Graubuenden Unterengadin Scuol Champatsch | 46.83 N | 10.26 E | 552 | 18,60 % |
| TLMF Lep 30311 | SAMN32653181 | *S. hochenwarthi* | Switzerland | Graubuenden Unterengadin Scuol Champatsch | 46.83 N | 10.26 E | 586 | 14,50 % |
| TLMF Lep 30312 | SAMN32653182 | *S. hochenwarthi* | Austria | Grossglockner Wallackhaus Kärnten |  |  | 542 | 18,31 % |
| TLMF Lep 30313 | SAMN32653183 | *S. hochenwarthi* | Austria | Grossglockner Wallackhaus Kärnten |  |  | 450 | 33,78 % |
| TLMF Lep 30314 | SAMN32653184 | *S. hochenwarthi* | Austria | Grossglockner Wallackhaus Kärnten |  |  | 555 | 18,11 % |
| MM27168 | SAMN32653185 | *S. hochenwarthi* | Finland |  |  |  | 569 | 17,09 % |
| MM27169 | SAMN32653186 | *S. hochenwarthi* | Finland |  |  |  | 567 | 16,75 % |
| MM27170 | SAMN32653187 | *S. hochenwarthi* | Finland |  |  |  | 565 | 17,08 % |
| MM27171 | SAMN32653188 | *S. hochenwarthi* | Finland |  |  |  | 548 | 17,98 % |
| MM27172 | SAMN32653189 | *S. hochenwarthi* | Finland |  |  |  | 496 | 24,91 % |
| MM27173 | SAMN32653190 | *S. hochenwarthi* | Finland |  |  |  | 528 | 21,15 % |
| MM27174 | SAMN32653191 | *S. hochenwarthi* | Finland |  |  |  | 557 | 17,05 % |
| TLMF Lep 04431 | SAMN32653192 | *X. speciosa* | Austria | Fohramoos ENE | 47.42 N | 9.81 E | 538 | 14,36 % |
| TLMF Lep 04432 | SAMN32653193 | *X. speciosa* | Austria | Fohramoos ENE | 47.42 N | 9.81 E | 565 | 10,10 % |
| TLMF Lep 02970 | SAMN32653194 | *X. speciosa* | Austria | Trojeralmtal, Ht. Trojeralm/ St. Jakob in Defereggen | 46.95 N | 12.31 E | 547 | 12,57 % |
| TLMF Lep 02969 | SAMN32653195 | *X. speciosa* | Italy | Pontebba/ Torbiera di Pramollo | 46.56 N | 13.29 E | 488 | 24,62 % |
| TLMF Lep 00183 | SAMN32653196 | *X. speciosa* | Austria |  | 47.419 N | 9.807 E | 539 | 14,82 % |
| TLMF Lep 30309 | SAMN32653197 | *X. speciosa* | Austria | Partenen, N Vermuntstausee | 46.94 N | 10.06 E | 550 | 11,79 % |
| TLMF Lep 02458 | SAMN32653198 | *X. speciosa* | Italy | Ritten/ Obergruenwald | 46.6 N | 11.44 E | 541 | 14,51 % |
| TLMF Lep 10975 | SAMN32653199 | *X. speciosa* | Austria | Hahntennjoch E | 47.29 N | 10.67 E | 534 | 16,57 % |
| TLMF Lep 20654 | SAMN32653200 | *X. speciosa* | Russia | 11 km NNW Aktash vill., Ajgulak Mts. Range | 50.42 N | 87.57 E | 563 | 10,17 % |
| TLMF Lep 20655 | SAMN32653201 | *X. speciosa* | Russia | 11 km NNW Aktash vill., Ajgulak Mts. Range | 50.42 N | 87.57 E | 576 | 8,37 % |
| TLMF Lep 30320 | SAMN32653202 | *X. speciosa* | Russia | Altai Republic Ulagan distr. Aktash vill. Ajgulak Mts. | 50.25 N | 87.34 E | 556 | 13,24 % |
| MM27368 | SAMN32653203 | *X. speciosa* | Finland | Kiiminki, Ostrobottnia ouluensis | 65.13 N | 25.78 E | 566 | 9,63 % |
| MM27414 | SAMN32653204 | *X. speciosa* | Finland | Korvasvaara | 65.96 N | 29.19 E | 558 | 9,44 % |
| MM27415 | SAMN32653205 | *X. speciosa* | Finland | Yppäri Leppikari | 64.46 N | 24.26 E | 539 | 12,47 % |
| MM27416 | SAMN32653206 | *X. speciosa* | Finland | Enontekijö, Lapponia enontekiensis | 68.38 N | 23.63 E | 542 | 12,83 % |
| MM27417 | SAMN32653207 | *X. speciosa* | Finland | Enontekijö, Lapponia enontekiensis | 68.38 N | 23.63 E | 579 | 7,59 % |
| TLMF Lep 22032 | SAMN32653208 | *A. glandon* | France | Col de la Bonette | 44.29 N | 6.90 E | 372 | 20,28 % |
| TLMF Lep 22033 | SAMN32653209 | *A. glandon* | France | Col de la Bonette | 44.29 N | 6.90 E | 391 | 16,78 % |
| TLMF Lep 22037 | SAMN32653210 | *A. glandon* | Switzerland | Arosa, Furggatobel | 46.77 N | 9.72 E | 367 | 23,31 % |
| TLMF Lep 22036 | SAMN32653211 | *A. glandon* | Switzerland | Arosa, Furggatobel | 46.77 N | 9.72 E | 405 | 14,01 % |
| TLMF Lep 22841 | SAMN32653212 | *A. glandon* | Italy | Franzenshoehe N/ Stilfserjochstrasse | 46.53 N | 10.46 E | 416 | 14,52 % |
| TLMF Lep 22842 | SAMN32653213 | *A. glandon* | Italy | Franzenshoehe N/ Stilfserjochstrasse | 46.53 N | 10.46 E | 420 | 11,51 % |
| TLMF Lep 30335 | SAMN32653214 | *A. glandon* | Switzerland | Graubuenden Unterengadin Scuol Champatsch | 46.83 N | 10.26 E | 422 | 10,76 % |
| TLMF Lep 30336 | SAMN32653215 | *A. glandon* | Switzerland | Graubuenden Fops/Fuorcla da Sagogn Nördl. Ilanz | 46.51 N | 9.10 E | 391 | 18,95 % |
| 07PROBE-00149 | SAMN32653216 | *A. glandon* | Canada | CAN: MB; Churchill | 58.77 N | 93.87 W | 410 | 12,68 % |
| 08BBLEP-04131 | SAMN32653217 | *A. glandon* | Canada | CAN: AB; Rocky Mountains; Banff National park | 51.06 N | 115.78 W | 410 | 13,23 % |
| 09BBELE-1273 | SAMN32653218 | *A. glandon* | Canada | CAN: Newfoundland: Terra Nova; Ochre's Hill | 48.51 N | 53.95 W | 404 | 14,07 % |
| 10BBCLP-0090 | SAMN32653219 | *A. glandon* | Canada | CAN: Saskatchewan; Prince Albert NP | 53.58 N | 106.05 W | 421 | 10,99 % |
| 10BBCLP-0183 | SAMN32653220 | *A. glandon* | Canada | CAN: British Columbia: Yoho NP, Takakkaw Falls | 51.50 N | 116.47 W | 418 | 11,07 % |
| BIOUG22492-G09 | SAMN32653221 | *A. glandon* | Canada | CAN: YT; Kluane National Park; Sheep Mountain | 61.01 N | 138.53 W | 423 | 9,58 % |
| BIOUG08958-A04 | SAMN32653222 | *A. glandon* | Canada | CAN: AB; Jasper NP | 53.19 N | 117.95 W | 424 | 10,99 % |
| TLMF Lep 06116 | SAMN32653223 | *A. caja* | Austria | Partenen, N Vermuntstausee | 46.94 N | 10.06 E | 838 | 14,92 % |
| TLMF Lep 20620 | SAMN32653224 | *A. caja* | Russia | 11 km NNW Aktash vill., Ajgulak Mts. Range | 50.42 N | 87.57 E | 863 | 8,87 % |
| TLMF Lep 20532 | SAMN32653225 | *A. caja* | Russia | 17 km NNE Kokorya vill., Chikhacheva Mts. Range, Talduair Mt., valley of Sajlyugem river | 50.02 N | 89.23 E | 824 | 18,52 % |
| TLMF Lep 20533 | SAMN32653226 | *A. caja* | Russia | 17 km NNE Kokorya vill., Chikhacheva Mts. Range, Talduair Mt., valley of Sajlyugem river | 50.02 N | 89.23 E | 897 | 8,16 % |
| TLMF Lep 30317 | SAMN32653227 | *A. caja* | Austria | Vorarlberg Lech-Zug | 47.10 N | 10.33 E | 898 | 6,65 % |
| TLMF Lep 30318 | SAMN32653228 | *A. caja* | Austria | Steiermark Wörschach Wörschacher Moos | 47.33 N | 14.10 E | 880 | 8,00 % |
| TLMF Lep 30319 | SAMN32653229 | *A. caja* | Austria | Torino PN Orsiera Rocciavre | 45.36 N | 7.42 E | 903 | 6,04 % |
| MM27153 | SAMN32653230 | *A. caja* | Finland | Kiiminki, Ostrobottnia ouluensis | 65.15 N | 25.84 E | 783 | 27,63 % |
| MM27154 | SAMN32653231 | *A. caja* | Finland | Otravaara | 61.89 N | 30.12 E | 896 | 6,67 % |
| MM27155 | SAMN32653232 | *A. caja* | Finland | Fiskars | 60.14 N | 23.49 E | 899 | 6,21 % |
| MM27156 | SAMN32653233 | *A. caja* | Finland | Vuorentaka | 60.35 N | 23.05 E | 864 | 11,03 % |
| MM27157 | SAMN32653234 | *A. caja* | Finland | Parikkala, Karelia ladogensis | 61.51 N | 29.55 E | 791 | 24,01 % |
| 2005-ONT-1199 | SAMN32653235 | *A. caja* | Canada | Ontario, Canada | 43.50 N | 80.10 W | 867 | 8,47 % |
| 2005-ONT-1163 | SAMN32653236 | *A. caja* | Canada | Puslinch, Ontario | 43.50 N | 80.10 W | 888 | 7,99 % |
| RWWA-0764 | SAMN32653237 | *A. caja* | USA | Bay Center; Edge of WillapaBay Wilson Residence, WA, USA | 46.62 N | 123.95 W | 867 | 9,22 % |
| RWWA-0812 | SAMN32653238 | *A. caja* | USA | Bay Center; Edge of WillapaBay Wilson Residence, WA, USA | 46.62 N | 123.95 W | 888 | 6,82 % |
| RWWA-0868 | SAMN32653239 | *A. caja* | USA | Bay Center; Edge of WillapaBay Wilson Residence, WA, USA | 46.62 N | 123.95 W | 865 | 8,44 % |
| MNBTT-3205 | SAMN32653240 | *A. caja* | Canada | York Co., NB McAdam | 45.58 N | 67.35 W | 883 | 7,73 % |
| BIOUG18201-F11 | SAMN32653241 | *A. caja* | Canada | CAN:Torngat mountains national park;saglek base camp | 58.45 N | 62.80 W | 903 | 6,33 % |
| TLMF Lep 17276 | SAMN32653242 | *C. sororiata* | Austria | Zirmbachalm W | 47.22 N | 11.03 E | 410 | 42,72 % |
| TLMF Lep 20717 | SAMN32653243 | *C. sororiata* | Russia | 11 km NNW Aktash vill., Ajgulak Mts. Range | 50.42 N | 87.57 E | 437 | 38,41 % |
| TLMF Lep 20718 | SAMN32653244 | *C. sororiata* | Russia | 11 km NNW Aktash vill., Ajgulak Mts. Range | 50.42 N | 87.57 E | 562 | 23,26 % |
| TLMF Lep 20719 | SAMN32653245 | *C. sororiata* | Russia | 11 km NNW Aktash vill., Ajgulak Mts. Range | 50.42 N | 87.57 E | 483 | 32,93 % |
| TLMF Lep 13731 | SAMN32653246 | *C. sororiata* | Austria | Loas zwischen Gamsstein und Kellerjoch-Hochleger | 47.30 N | 11.76 E | 552 | 23,18 % |
| TLMF Lep 02795 | SAMN32653247 | *C. sororiata* | Austria | Karn.A., Untertilliach, Winklertal | 46.68 N | 12.65 E | 504 | 30,28 % |
| TLMF Lep 02796 | SAMN32653248 | *C. sororiata* | Austria | Karn.A., Untertilliach, Winklertal | 46.68 N | 12.65 E | 518 | 28,05 % |
| TLMF Lep 12550 | SAMN32653249 | *C. sororiata* | Austria | Partenen, Zeinissee | 46.98 N | 10.12 E | 517 | 26,78 % |
| MM27370 | SAMN32653250 | *C. sororiata* | Finland | Pyöriäsuo | 64.88 N | 25.64 E | 613 | 17,20 % |
| MM27371 | SAMN32653251 | *C. sororiata* | Finland | Pyöriäsuo | 64.88 N | 25.64 E | 567 | 21,62 % |
| MM27372 | SAMN32653252 | *C. sororiata* | Finland | Vuorentaka | 60.35 N | 23.05 E | 578 | 19,89 % |
| MM27373 | SAMN32653253 | *C. sororiata* | Finland | Kylmäkoski, Tavastia australis | 61.13 N | 23.57 E | 530 | 26,36 % |
| MM27374 | SAMN32653255 | *C. sororiata* | Finland | Finnmossen | 62.76 N | 17.88 E | 522 | 26,57 % |
| MM27375 | SAMN32653254 | *C. sororiata* | Finland | Kylmäkoski, Tavastia australis | 61.13 N | 23.57 E | 492 | 29,76 % |
| 04HBL003625 | SAMN32653256 | *C. sororiata* | Canada | CAN: Churchill, Manitoba | 58.70 N | 93.80 W | 469 | 34,11 % |
| 06-PROBE-2668 | SAMN32653257 | *C. sororiata* | Canada | CAN: MB: Churchill | 58.44 N | 93.49 W | 543 | 24,61 % |
| BIOUG11511-H02 | SAMN32653258 | *C. sororiata* | Canada | CAN: NL; Torngat Mountains National Park; Ivitak valley | 58.95 N | 63.68 W | 627 | 14,72 % |
| BIOUG11511-H03 | SAMN32653259 | *C. sororiata* | Canada | CAN: NL; Torngat Mountains National Park; Ivitak valley | 58.95 N | 63.68 W | 641 | 13,79 % |
| BIOUG55218-F10 | SAMN32653260 | *C. sororiata* | Canada | CAN: YT; Dàadzàii Vàn Territorial Park | 67.52 N | 136.50 W | 621 | 15,49 % |
| BIOUG17479-G01 | SAMN32653261 | *C. sororiata* | Canada |  |  |  | 588 | 18,34 % |
| TLMF Lep 17436 | SAMN32653262 | *C. tullia* | Austria | Dollinger/ Imst NE | 47.28 N | 10.79 E | 344 | 47,16 % |
| TLMF Lep 17435 | SAMN32653263 | *C. tullia* | Austria | Schwendt, Enzianwiese | 47.63 N | 12.38 E | 364 | 42,03 % |
| TLMF Lep 19575 | SAMN32653264 | *C. tullia* | Austria | Spechtensee | 47.56 N | 14.09 E | 339 | 45,27 % |
| TLMF Lep 19576 | SAMN32653265 | *C. tullia* | Austria | Spechtensee | 47.56 N | 14.09 E | 283 | 59,83 % |
| TLMF Lep 10011 | SAMN32653266 | *C. tullia* | Austria | Im Moos/ Bizau W | 47.37 N | 9.91 E | 421 | 34,14 % |
| MM27158 | SAMN32653267 | *C. tullia* | Finland | Kiiminki, Ostrobottnia ouluensis | 65.07 N | 25.95 E | 500 | 24,99 % |
| MM23885 | SAMN32653268 | *C. tullia* | Finland |  | 64.43 N | 28.19 E | 392 | 37,14 % |
| MM23886 | SAMN32653269 | *C. tullia* | Finland |  | 63.91 N | 29.84 E | 390 | 38,53 % |
| MM27159 | SAMN32653270 | *C. tullia* | Finland |  | 65.11 N | 25.62 E | 373 | 41,01 % |
| MM27160 | SAMN32653271 | *C. tullia* | Finland |  | 64.75 N | 25.01 E | 400 | 36,76 % |
| MM27161 | SAMN32653272 | *C. tullia* | Finland |  | 60.18 N | 23.02 E | 528 | 21,01 % |
| 09BBLEP-04021 | SAMN32653273 | *C. tullia* | USA | USA: New Mexico: San Migual; Jack's creek campground | 35.84 N | 105.66 W | 439 | 31,24 % |
| 09BBLEP-04288 | SAMN32653274 | *C. tullia* | USA | USA: Colorado: El PasoCo; Golden Eagle ranch | 38.71 N | 104.84 W | 477 | 25,41 % |
| 09BBELE-1280 | SAMN32653275 | *C. tullia* | Canada | CAN: Newfoundland: Terra Nova; Ochre's Hill | 48.51 N | 53.95 W | 472 | 26,04 % |
| 10BBCLP-0098 | SAMN32653276 | *C. tullia* | Canada | CAN: British Columbia: Yoho NP | 51.42 N | 116.43 W | 504 | 23,04 % |
| BIOUG01573-E01 | SAMN32653277 | *C. tullia* | Canada | CAN: Ontario | 44.35 N | 76.89 W | 517 | 21,35 % |
| CHARS00315-F08 | SAMN32653278 | *C. tullia* | Canada | CAN: Nunavut; Kitikmeot; Kugluktuk Kugluk | 67.74 N | 115.37 W | 513 | 21,29 % |
| TLMF Lep 20645 | SAMN32653279 | *M. brunneata* | Russia | 11 km NNW Aktash vill., Ajgulak Mts. Range | 50.42 N | 87.57 E | 466 | 23,94 % |
| TLMF Lep 20711 | SAMN32653280 | *M. brunneata* | Russia | 11 km NNW Aktash vill., Ajgulak Mts. Range | 50.42 N | 87.57 E | 425 | 29,27 % |
| TLMF Lep 20644 | SAMN32653281 | *M. brunneata* | Russia | 11 km NNW Aktash vill., Ajgulak Mts. Range | 50.42 N | 87.57 E | 457 | 25,91 % |
| TLMF Lep 17068 | SAMN32653282 | *M. brunneata* | Austria | Bielerhoehe SE: Bieltal | 46.91 N | 10.12 E | 430 | 28,09 % |
| TLMF Lep 18713 | SAMN32653283 | *M. brunneata* | Austria | Fliess NE/ Pillermoor | 47.12 N | 10.67 E | 446 | 25,59 % |
| TLMF Lep 30327 | SAMN32653284 | *M. brunneata* | Italy | Pfitscherjoch | 46.59 N | 11.40 E | 447 | 24,86 % |
| TLMF Lep 30315 | SAMN32653285 | *M. brunneata* | Austria | Nordtirol Kuehtai | 47.13 N | 11.02 E | 455 | 25,00 % |
| TLMF Lep 30316 | SAMN32653286 | *M. brunneata* | Austria | Nordtirol Zillertal Aschtau Riedaste | 47.16 N | 11.52 E | 461 | 24,11 % |
| MM27162 | SAMN32653287 | *M. brunneata* | Finland |  | 65.12 N | 25.83 E | 524 | 17,07 % |
| MM27163 | SAMN32653289 | *M. brunneata* | Finland |  | 64.88 N | 25.64 E | 376 | 37,04 % |
| MM27164 | SAMN32653288 | *M. brunneata* | Finland |  | 65.12 N | 25.83 E | 329 | 43,74 % |
| MM27165 | SAMN32653290 | *M. brunneata* | Finland |  | 63.91 N | 29.84 E | 404 | 32,19 % |
| MM27166 | SAMN32653291 | *M. brunneata* | Finland |  | 59.81 N | 23.21 E | 360 | 37,95 % |
| 08BBLEP-02562 | SAMN32653292 | *M. brunneata* | Canada | CAN: AB south; Waterton Lakes Nat. Park | 49.09 N | 113.97 W | 476 | 21,78 % |
| 10BBCLP-0565 | SAMN32653293 | *M. brunneata* | Canada | CAN: British Columbia: Kootenay NP | 50.76 N | 115.94 W | 458 | 23,26 % |
| BIOUG06593-B05 | SAMN32653294 | *M. brunneata* | Canada | CAN: NT; Wood Buffalo National Park | 59.56 N | 112.26 W | 488 | 20,16 % |
| BIOUG06593-G01 | SAMN32653295 | *M. brunneata* | Canada | CAN: NT; Wood Buffalo National Park | 59.56 N | 112.26 W | 481 | 20,47 % |
| BIOUG15746-B02 | SAMN32653296 | *M. brunneata* | Canada | CAN: NT; Nahanni National Park; Nailicho (virginia falls) | 61.61 N | 125.76 W | 542 | 14,34 % |
| BIOUG45338-C06 | SAMN32653297 | *M. brunneata* | Canada | CAN: YT; Whitehorse; Crestview Springs | 60.79 N | 135.19 W | 520 | 16,71 % |
| BIOUG45338-E10 | SAMN32653298 | *M. brunneata* | Canada | CAN: YT; Whitehorse; Crestview Springs | 60.79 N | 135.19 W | 505 | 17,61 % |
| TLMF Lep 00747 | SAMN32653299 | *X. lorezi* | Austria | Kleiner Trieb E | 47.0 N | 13.07 E | 580 | 17,63 % |
| TLMF Lep 30323 | SAMN32653300 | *X. lorezi* | Austria | Vorarlberg Formarinsee | 47.103 N | 10.00 E | 605 | 9,97 % |
| TLMF Lep 30324 | SAMN32653301 | *X. lorezi* | Austria | Vorarlberg Formarinsee | 47.101 N | 10.01 E | 602 | 9,52 % |
| TLMF Lep 30325 | SAMN32653302 | *X. lorezi* | Austria | Vorarlberg Formarinsee | 47.1014 N | 10.012 E | 625 | 7,67 % |
| TLMF Lep 30326 | SAMN32653303 | *X. lorezi* | Germany | Nebelhorn |  |  | 583 | 13,70 % |
| MM08265 | SAMN32653304 | *X. lorezi* | Finland |  | 69.04 N | 20.82 E | 580 | 12,43 % |
| MM15896 | SAMN32653305 | *X. lorezi* | Finland |  | 69.05 N | 20.85 E | 496 | 29,94 % |
| MM26177 | SAMN32653308 | *X. lorezi* | Austria |  | 47.17 N | 10.01 E | 617 | 8,19 % |
| MM26178 | SAMN32653309 | *X. lorezi* | Finland |  | 69.42 N | 26.12 E | 612 | 9,56 % |
| MM26179 | SAMN32653306 | *X. lorezi* | Finland |  | 69.05 N | 20.85 E | 638 | 6,16 % |
| MM26180 | SAMN32653307 | *X. lorezi* | Finland |  | 69.05 N | 20.85 E | 637 | 5,99 % |
| TLMF Lep 30363 | SAMN32653310 | *X. lorezi* |  |  |  |  | 591 | 11,95 % |
| TLMF Lep 30366 | SAMN32653311 | *X. lorezi* |  |  |  |  | 636 | 6,00 % |
| CNCNoctoidea12387 | SAMN32653312 | *X. lorezi* | North America |  |  |  | 209 | 75,27 % |
| CNC LEP00052808 | SAMN32653313 | *X. lorezi* | North America |  |  |  | 632 | 7,02 % |

Following command was used to extract the COI gene region from target enrichment assemblies -

‘exonerate --model est2genome -n 1 --ryo ">%ti (%tab - %tae)\n%tas\n” --showvulgar no --showalignment no COI_Consensus.fasta assembly.fas > COI.out)

Barcoding laboratory protocol –

The COI was amplified for using the primers HybLCO and HybHCO, and PCR was conducted under the following conditions: 95c 5min, (40x 95c 30s, 50c 30s, 72c 2min) and 72c 2min.

PCR purification was done using the Exo-sap purification method according to the following instructions - PCR product 4µl, Fast AP 0.5µl, FA buffer 0.8, EXO I 0.1µl and water 4.6µl. The final volume of the purified product was 10µl.

5µl of purified PCR product and 5µl of 5µm primer were sent for sequencing.

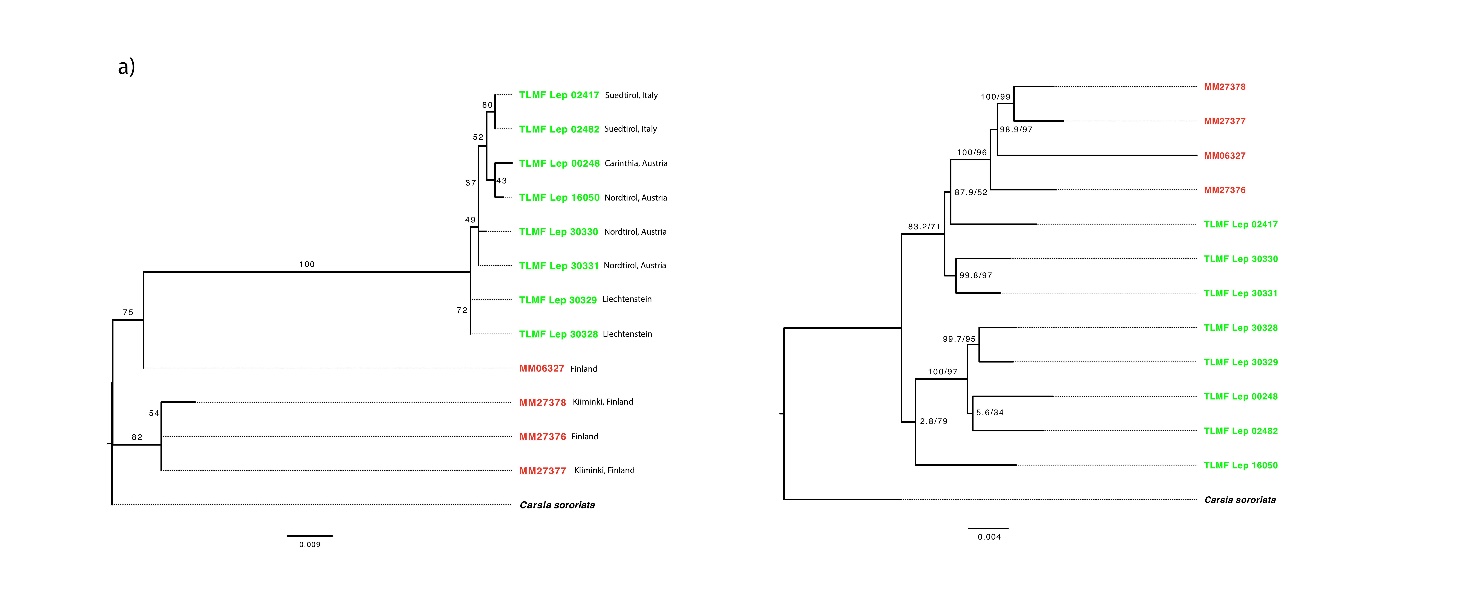

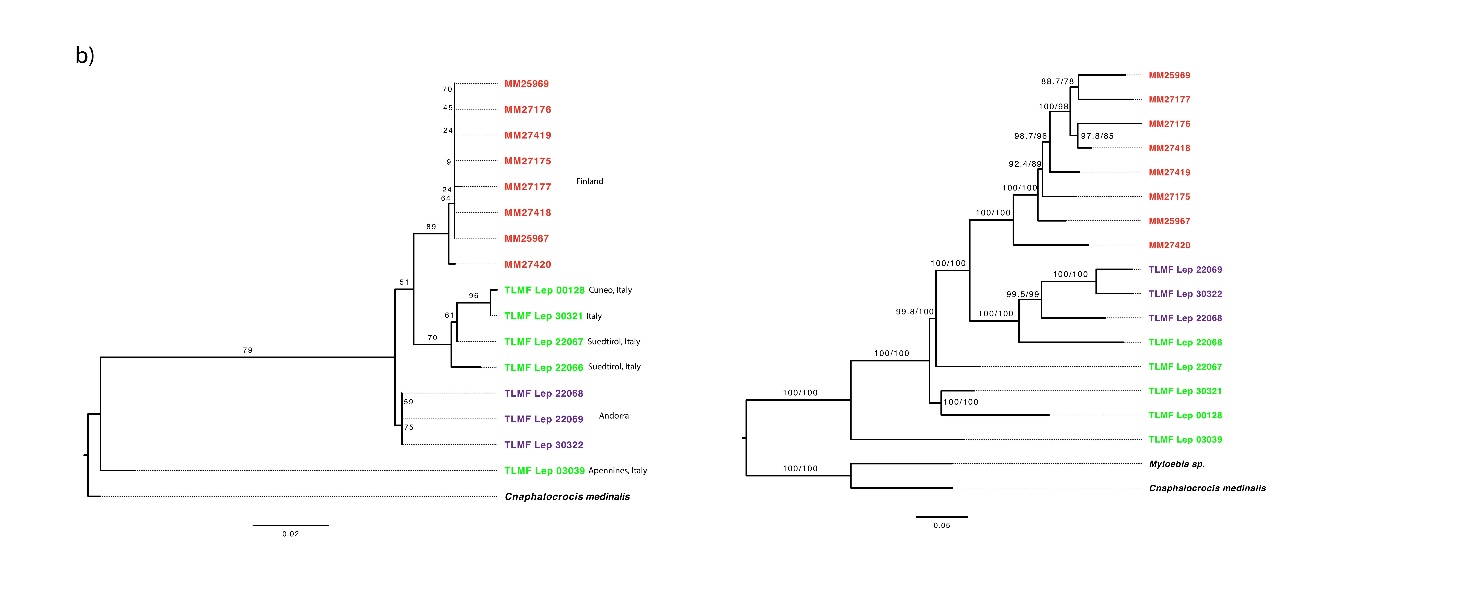

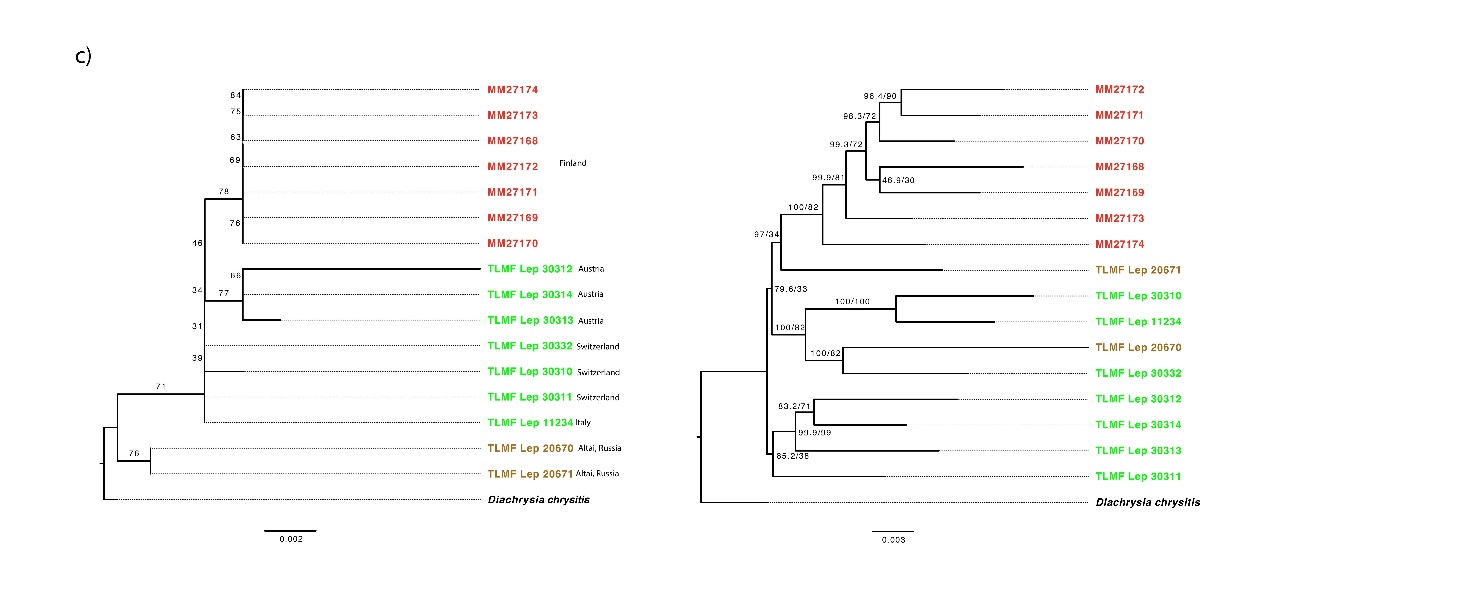

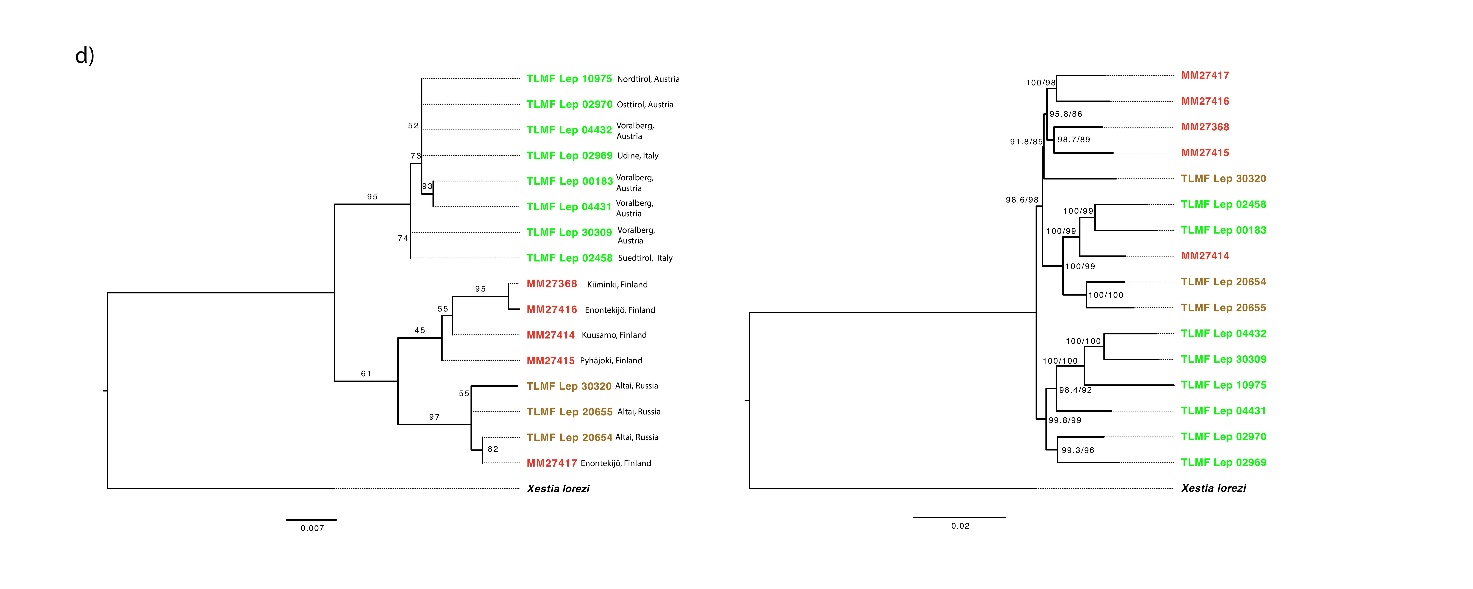

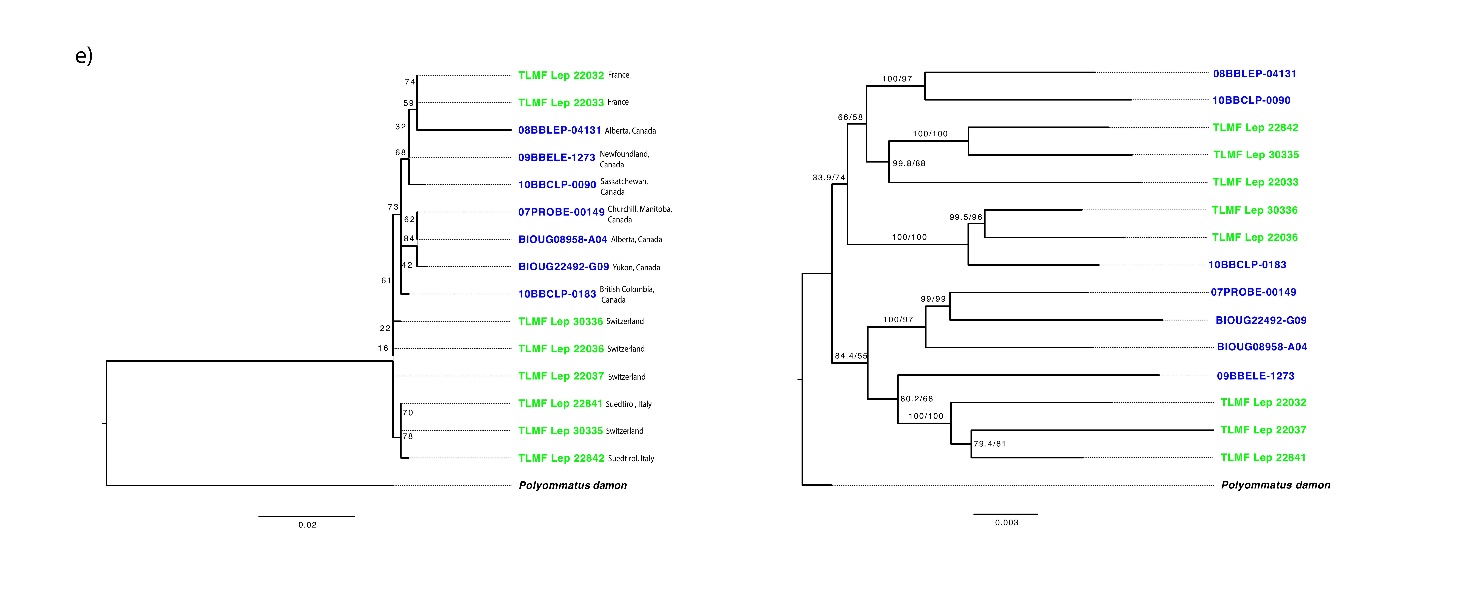

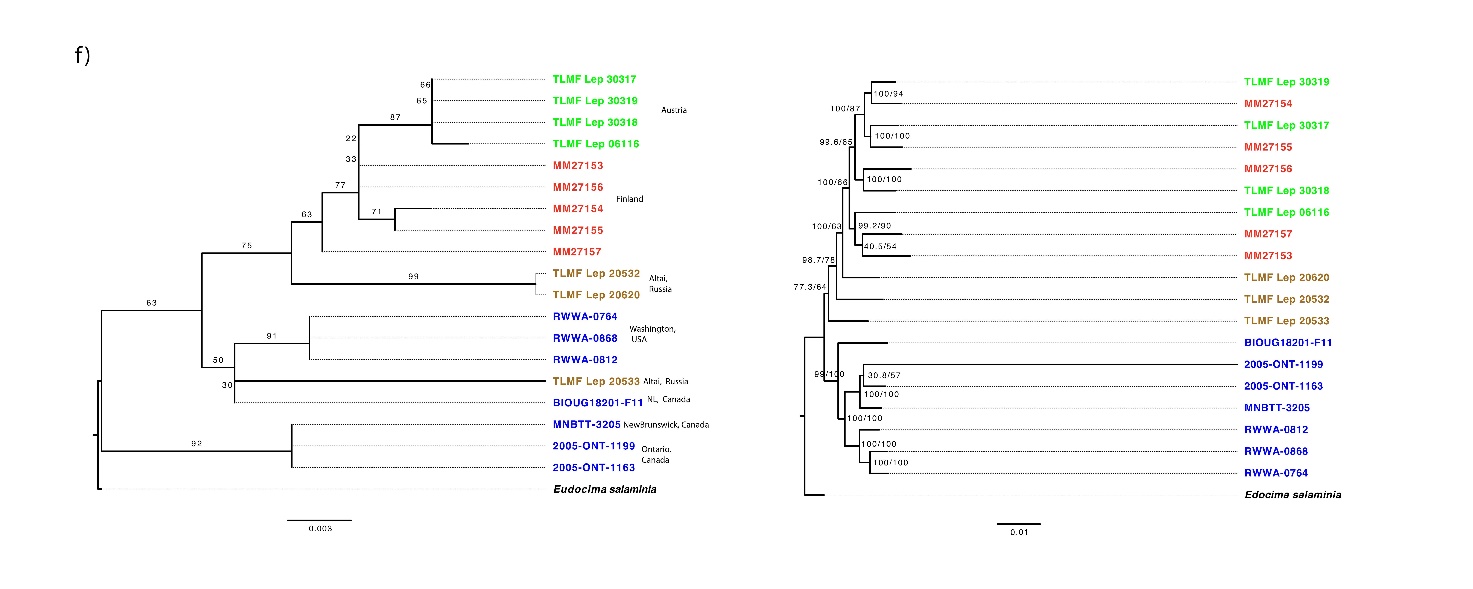

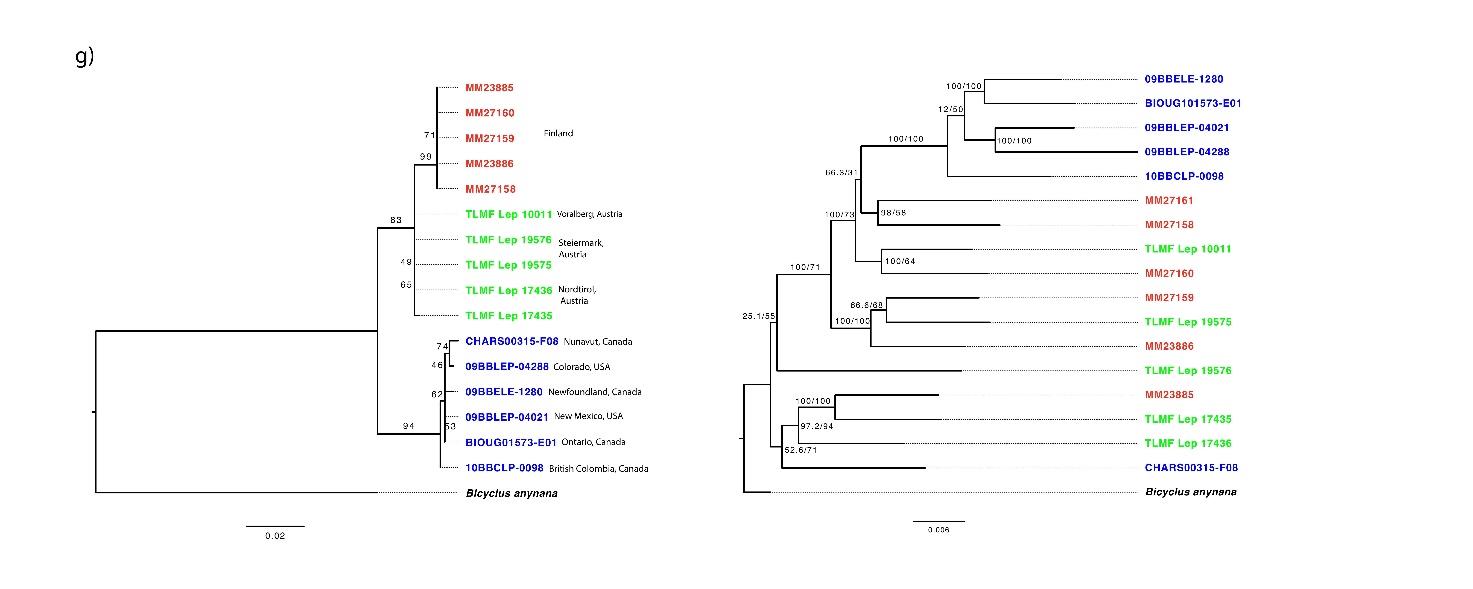

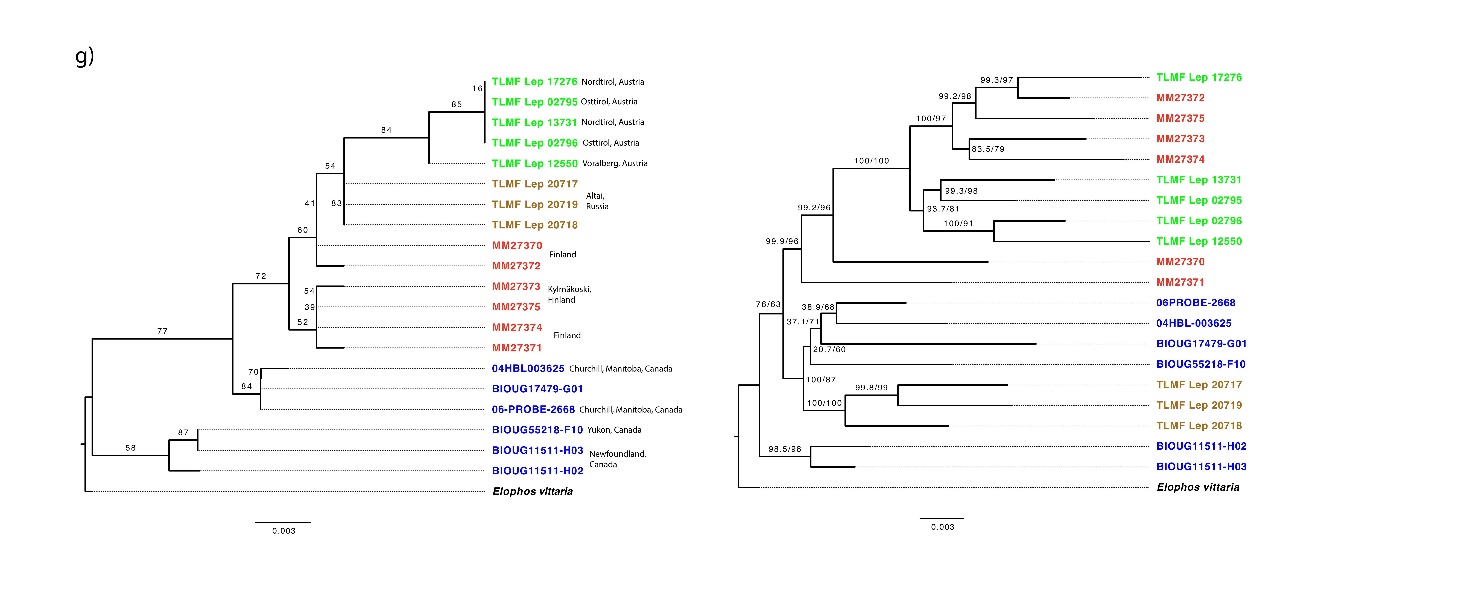

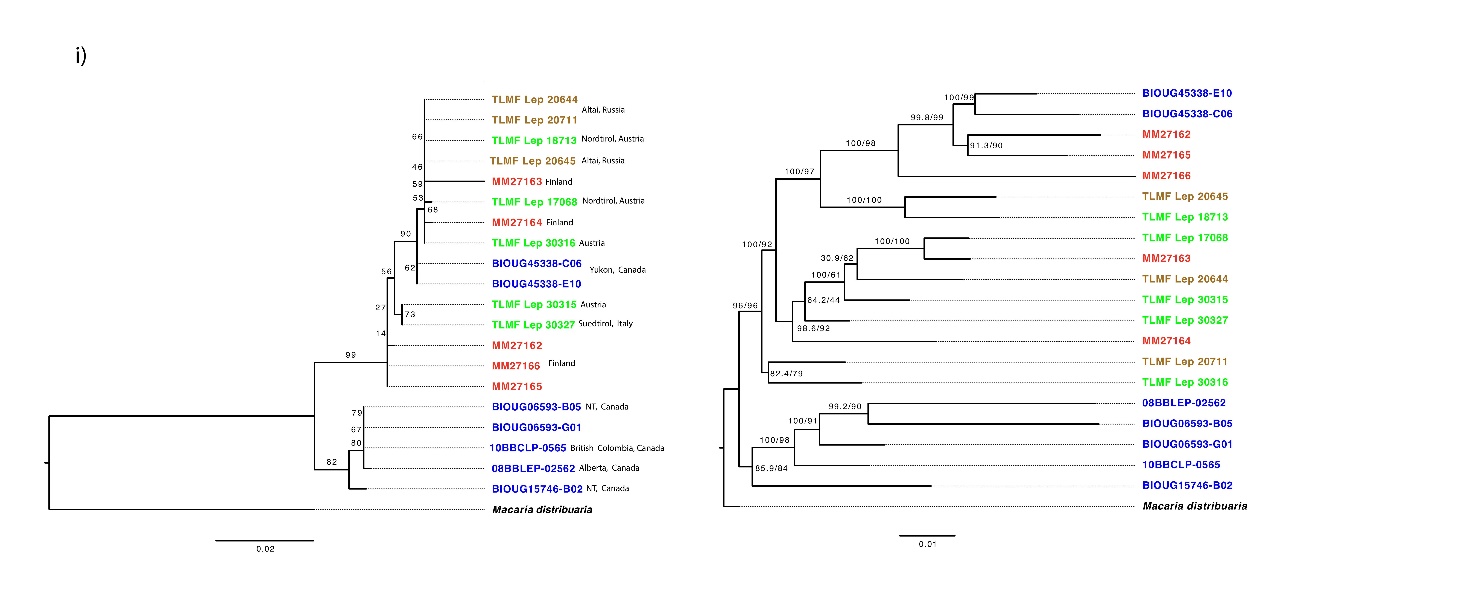

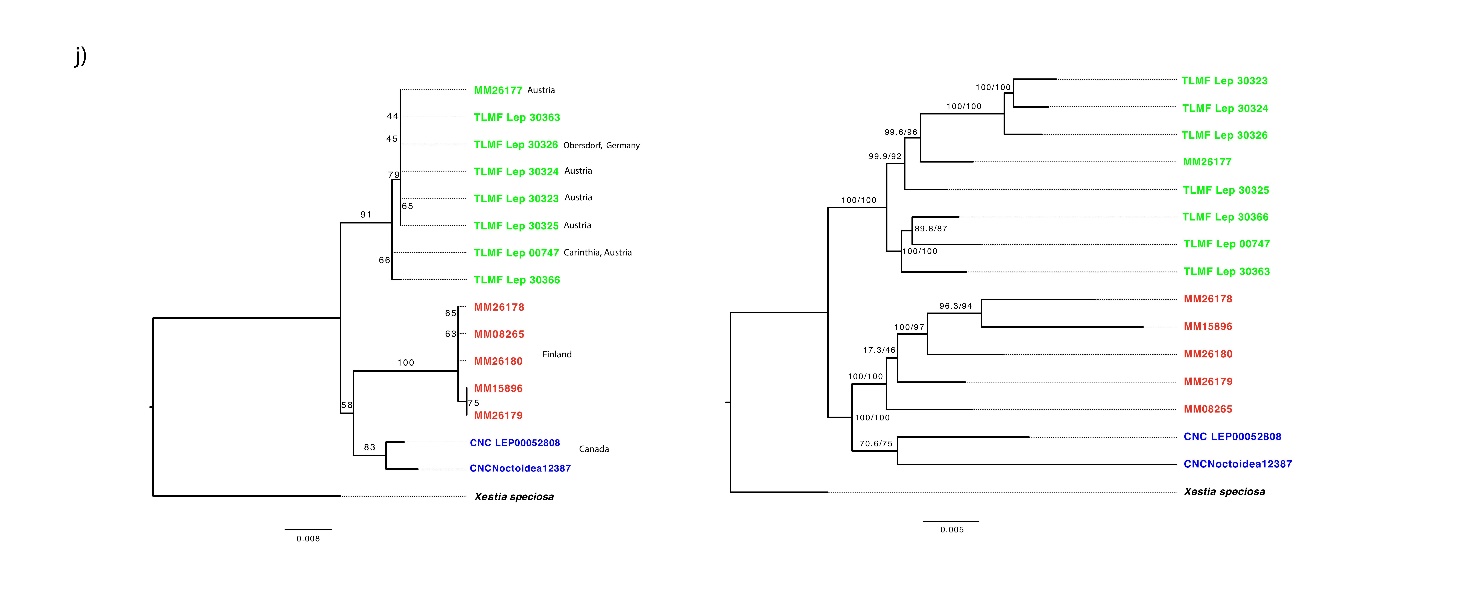
**Figure S1**: Maximum likelihood trees using barcode (on the left) and target enrichment (on the right) data for a) *E. vittaria* b) *E. sudetica* c) *S. hochenwarthi* d) *X. speciosa* e) *A. glandon* f) *A. caja* g) *C. sororiata* h) *C. tullia* i) *M. brunneata* j) *X. lorezi.* For barcode trees, numbers on the branches indicate ultrafast bootstrap support values based on 1000 replicates. For target enrichment ML trees, numbers on the branches/at the nodes indicates-aLRT/ultrafast bootstrap support values based on 1000 replicates.

Isolation by Distance plots -Fig S2

i. *X. speciosa*

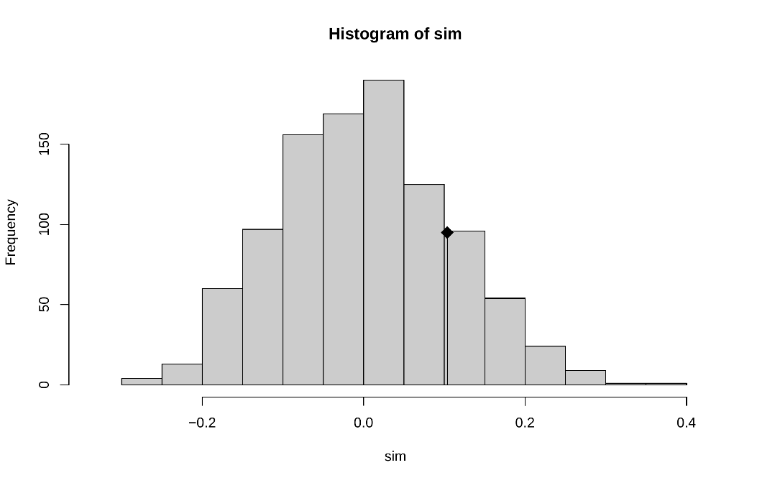

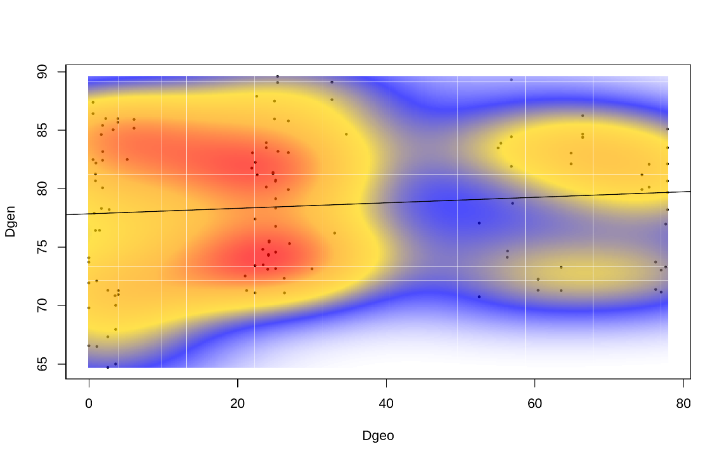

ii. *A. caja*

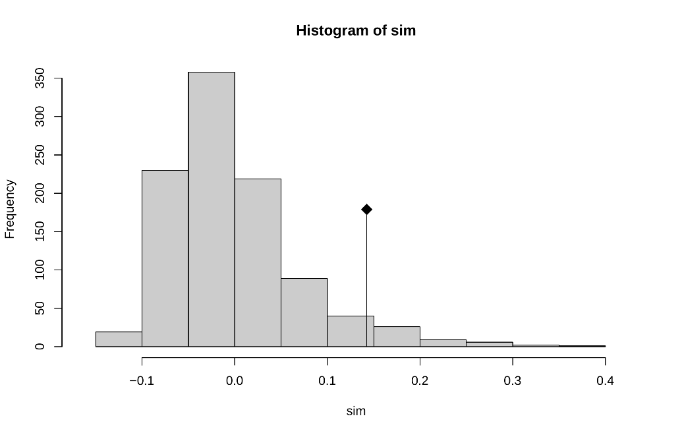

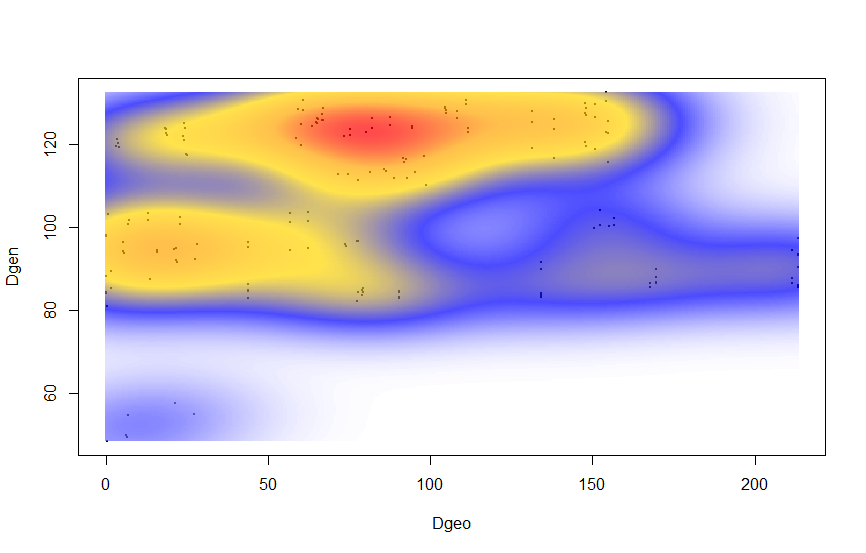

iii. *C. sororiata*

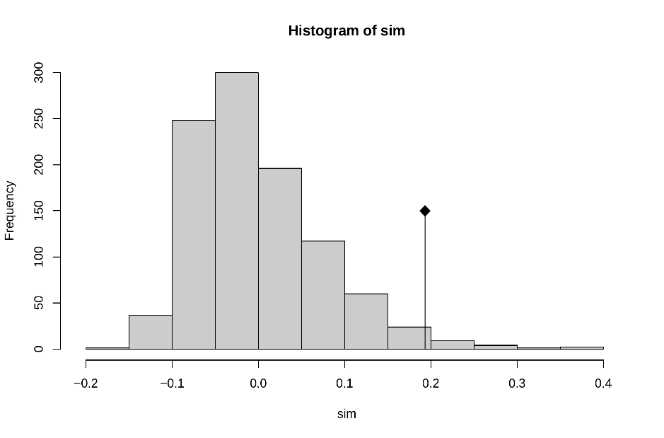

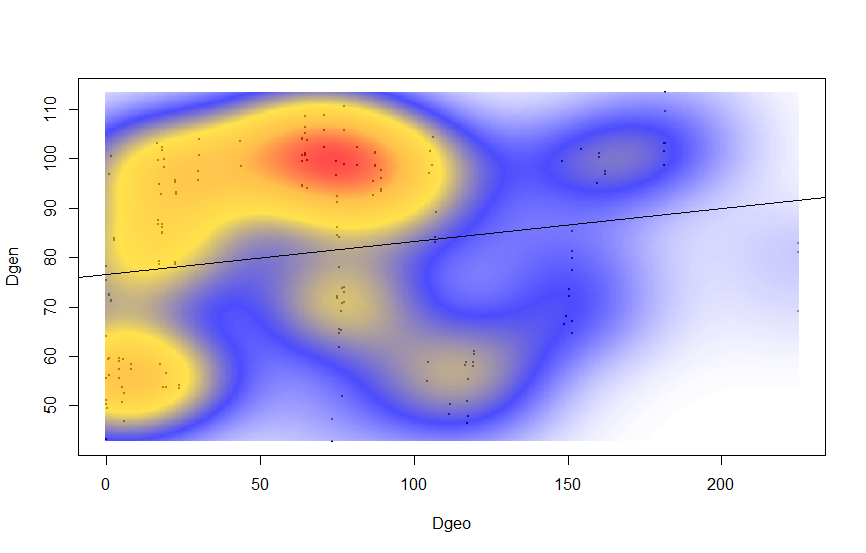

iv. *C. tullia*

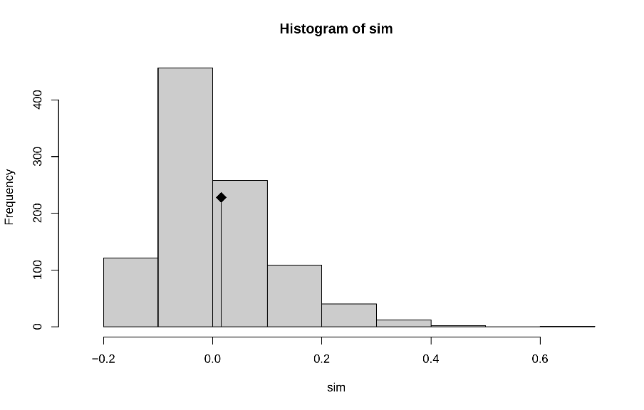

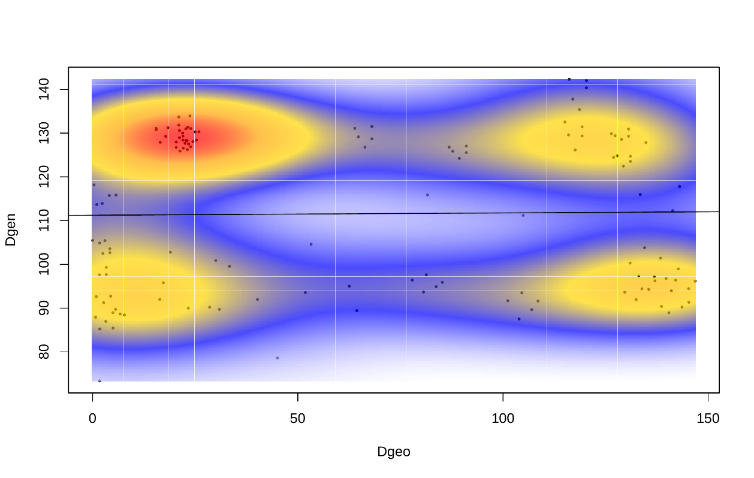

v. *M. brunneata*

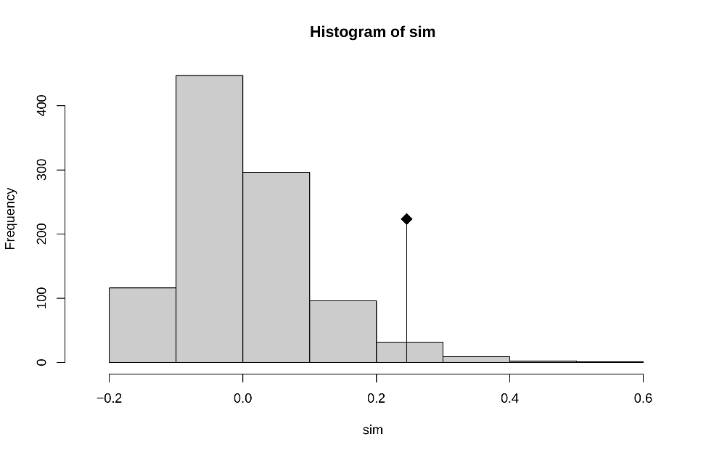

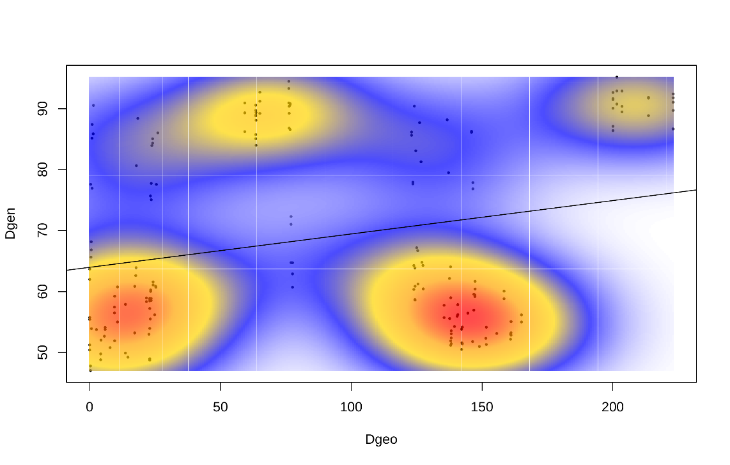

**Figure S2**: The histograms represent the permuted values (in absence of a spatial structure) of a correlation between two distance matrices – Edwards’ distance and Euclidean geographic distance. The original value of a correlation is represented by the dot, which represents a significant spatial structure if it is out of the reference distribution. On the right, the scatterplot of the correlation matrix is shown where local density is measured using 2-dimentional kernel density estimation.

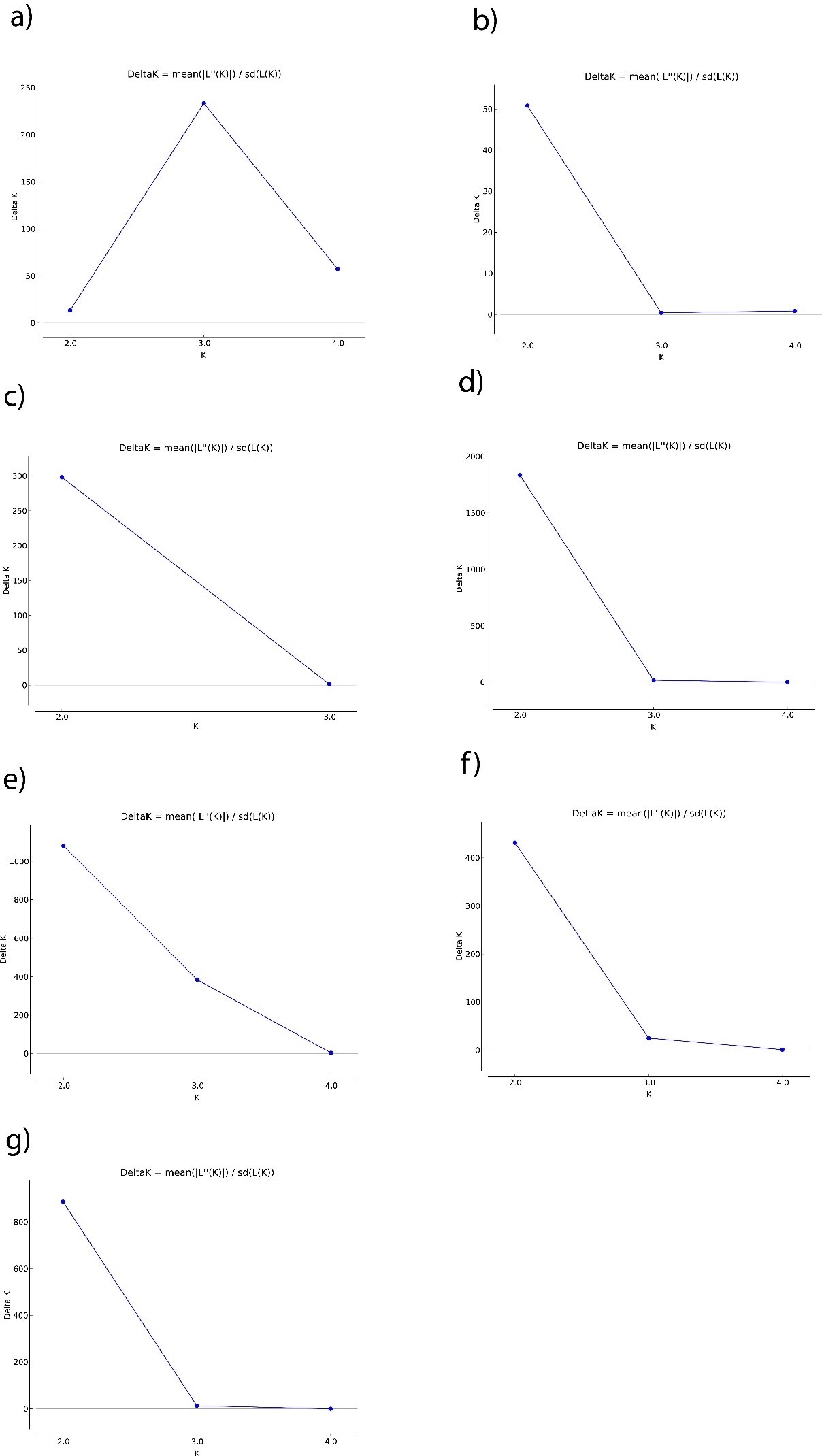

**Figure S3**: STRUCTURE ΔK plots for a) *E. sudetica* b) *S. hochenwarthi* c) *X. speciosa* d) *A. caja* e) *C. sororiata* f) *C. tullia* g) *M. brunneata*

**Tr2 ANALYSES**

1) ***E. vittaria***

null – All populations belong to same species – 23838.80

Model 1 – Fennoscandian and Alps populations as separate species – 213.20

2) ***E. sudetica***

null - All populations belong to same species – 64194.20

Model 1 – Fennoscandian population as separate species, Alps + Andorra same species – 2240.32

Model 2 – All three populations (FEN, Alps, Andorra separate species) – 819.14

Model 3 – Alps + FEN same species, Andorra population as separate species – 10496.38

3) ***S. hochenwarthi***

null – 59196.79

Model 1 – FEN as separate species, Alps+Altai same species – 801.28

Model 2 – All three populations are separate species – 793.43

Model 3 – Altai population separate species, Alps+FEN same species – 3808.87

4) ***X. speciosa***

null – 62599.62

Model 1 – FEN population separate species, Alps+Altai same species – 925.56

Model 2 – All three populations separate species – 857.18

Model 3 – Alps population as separate species, FEN+Altai same species – 762.82

5) ***A. glandon***

null – 34274.92

Model 1 – NA and Alps populations are separate species – 1025.90

6) ***A. caja***

null – 108105.52

Model 1 – NA population separate species, Alps+Altai+FEN same species – 2252.0

Model 2 – All populations are separate species – 1978.17

7) ***C. sororiata***

null – 89435.43

Model 1 – NA population separate species, Alps+Altai+FEN same species – 5378.51

Model 2 – NA+Altai same species, Alps+FEN same species – 1345.31

Model 3 – All 4 populations are separate species – 792.36

8) ***C. tullia***

null – 50673.59

Model 1 – NA population separate species, FEN+Alps same species – 574.14

Model 2 – All three populations are separate species – 995.51

9) ***M. brunneata***

null – 38611.72

Model 1 – NA population as separate species, FEN+Alps+Altai same species – 1081.48

Model 2 - All populations are separate species – 1938.41

Model 3 – NA population except for Yukon individual as separate species, FEN+Alps+Altai+Yukon individuals same species – 362.56

10) ***X. lorezi***

null – 68511.37

Model 1 – All three populations are separate species – 354.31

Model 2 – NA+FEN same species, Alps population is a separate species – 638.39
